# Quantitative Structure-Mutation-Activity Relationship Tests (QSMART) Model for Protein Kinase Inhibitor Response Prediction

**DOI:** 10.1101/868067

**Authors:** Liang-Chin Huang, Wayland Yeung, Ye Wang, Huimin Cheng, Aarya Venkat, Sheng Li, Ping Ma, Khaled Rasheed, Natarajan Kannan

## Abstract

Predicting drug sensitivity profiles from genotypes is a major challenge in personalized medicine. Machine learning and deep neural network methods have shown promise in addressing this challenge, but the “black-box” nature of these methods precludes a mechanistic understanding of how and which genomic and proteomic features contribute to the observed drug sensitivity profiles. Here we provide a combination of statistical and neural network framework that not only estimates drug IC_50_ in cancer cell lines with high accuracy (R^2^ = 0.861 and RMSE = 0.818) but also identifies features contributing to the accuracy, thereby enhancing explainability. Our framework, termed QSMART, uses a multi-component approach that includes (1) collecting drug fingerprints, cancer cell line’s multi-omics features, and drug responses, (2) testing the statistical significance of interaction terms, (3) selecting features by Lasso with Bayesian information criterion, and (4) using neural networks to predict drug response. We evaluate the contribution of each of these components and use a case study to explain the biological relevance of several selected features to protein kinase inhibitor response in non-small cell lung cancer cells. Specifically, we illustrate how interaction terms that capture associations between drugs and mutant kinases quantitatively contribute to the response of two EGFR inhibitors (afatinib and lapatinib) in non-small cell lung cancer cells. Although we have tested QSMART on protein kinase inhibitors, it can be extended across the proteome to investigate the complex relationships connecting genotypes and drug sensitivity profiles.

## Introduction

Protein kinases are a class of signaling proteins, greatly valued as therapeutic targets for their key roles in human diseases, such as cancer [1]. For decades, chemotherapy has served as part of a standard set of cancer treatments; however, the resistance of cancer cells to chemotherapy is still a major clinical challenge [2]. Mutations in protein kinase are known to play important roles not only in drug resistance [3] but also in drug sensitivity [4]. Depending on the structural location, mutations can have varying impacts on drug sensitivity. For example, non-small cell lung cancer (NSCLC) cells harboring either the EGFR T790M or L858R mutation respectively leads to resistance or hypersensitivity to the cancer drug gefitinib [5, 6], while those with EGFR T790M/L858R double mutant are only resistant to gefitinib [7]. As mutations impact the efficacy of different cancer drugs, there is a need to incorporate structural knowledge in drug response prediction methods.

To facilitate the understanding of the molecular mechanisms that cause drug sensitivity and drug resistance in cancer cells, the Genomics of Drug Sensitivity in Cancer (GDSC) Project [8] recently screened the drug responses of 266 anti-cancer drugs against ∼1,000 human cancer cell lines and provided the largest publicly available drug response dataset. Moreover, to broaden the pharmacologic annotation for human cancers, the Cancer Cell Line Encyclopedia [9] (CCLE) provided pharmacologic profiles for 24 drugs across 504 cancer cell lines. By utilizing these datasets, several prediction models were built to pursue a more accurate drug response estimation by different types of approaches, from traditional statistical models, network-based models, to the recent machine learning methods and state-of-the-art neural networks (Table 1). These approaches include (1) statistical models: MANOVA [10] and generalized linear models (regularization: ridge [11–14], elastic net [11–13, 15], Lasso [11–13], and mixture [16]), (2) network-based models [17–24], (3) random forests [25, 26], (4) support vector machine (SVM) [21, 27, 28] and other kernelized methods [29–31], and (5) neural networks: artificial neural network (ANN) [32], convolutional neural network (CNN) [33–35], recurrent neural network (RNN) [35], and other deep neural networks (DNN) [36, 37].

**Table 1.** Current drug response prediction approaches.

| Date | Author | Best model (Compared models) | Cancer cell line feature | Drug response |  | Validation | Performance |
| --- | --- | --- | --- | --- | --- | --- | --- |
|  |  |  |  | GDSC | CCLE |  |  |
| 2013.04.30 | Menden et al. [32] | ANN (RF) | MUT, CNV | ✓ | | 8-fold CV | $R^2 = 0.72$ |
| 2014.03.03 | Geeleher et al. [11] | GLM | EXP | ✓ |  | LOOCV | AUC = 0.81 |
| 2015.01.01 | Jang et al. [12] | GLM (PLS, SVM, PCA, RF) | MUT, EXP, CNV, CLS | ✓ | ✓ | 5-fold CV | $r = \sim 0.5$ |
| 2015.06.30 | Dong et al. [27] | SVM | EXP | | ✓ | 10-fold CV | Accuracy = $\sim 0.8$ |
| 2015.09.29 | Zhang et al. [17] | Network (EN) | EXP | ✓ | ✓ | LOOCV | $r = 0.6$ |
| 2016.03.31 | Gupta et al. [28] | SVM | MUT, EXP, CNV | | ✓ | LOOCV | $r = 0.78$ |
| 2016.09.01 | Ammad-ud-din et al. [29] | Kernel (GLM) | PWY | ✓ | | 5-fold CV | $\rho = \sim 0.22$ |
| 2016.12.28 | Nguyen et al. [10] | MANOVA (RF) | EXP | ✓ |  | 10-fold CV | MCC = 0.18 |
| 2017.01.09 | Stanfield et al. [18] | Network (Kernel) | MUT, PPI | ✓ | ✓ | LOOCV | AUC = 0.881 |
| 2017.07.15 | Ammad-ud-din et al. [13] | GLM (PLS, SGL, RF, SVM) | EXP, PWY | ✓ | | LOOCV | $\rho = 0.375$ |
| 2017.08.28 | Geeleher et al. [14] | Ridge | EXP | ✓ | | 10-fold CV | $\rho = 0.48$ |
| 2017.09.12 | Rahman et al. [25] | RF | EXP | ✓ | ✓ | 3-fold CV | AUC = $\sim 0.3$ |
| 2017.11.13 | Ding et al. [36] | DNN (EN, SVM) | MUT, EXP, CNV | ✓ | ✓ | 25-fold CV | AUC = 0.87 |
| 2018.03.08 | He et al. [30] | Kernel (EN, Ridge, RF) | EXP | ✓ | | 3-fold CV | Precision = $\sim 0.35$ |
| 2018.06.11 | Chang et al. [33] | CNN (RF, SVM) | SNP | ✓ | | 5% leave-out | $R^2 = 0.843$ |
| 2018.07.01 | Cichonska et al. [31] | Kernel | SNP, MET, EXP, CNV | ✓ | | 10-fold CV | $r = 0.858$ |
| 2018.09.14 | Le et al. [19] | Network (Kernel) | MUT, EXP | ✓ | ✓ | 5-fold CV | $r = 0.804$ |
| 2018.09.14 | Juan-Blanco et al. [20] | Network | MUT, EXP | | ✓ | LOOCV | AUC = $\sim 0.72$ |
| 2018.10.10 | Yang et al. [21] | Network + SVM (Kernel) | MUT, MET, CNV, PPI | ✓ |  | 5-fold CV | AUC = 0.788 |
| 2018.12.07 | Liu et al. [22] | Network | EXP | ✓ | ✓ | 10-fold CV | $r = 0.73$ |
| 2019.01.22 | Wei et al. [23] | Network | EXP | ✓ | ✓ | LOOCV | $r = 0.63$ |
| 2019.01.31 | Wang et al. [15] | EN | EXP, PWY | | ✓ | 10-fold CV | MSE = $\sim 2.8$ |
| 2019.01.31 | Chiu et al. [37] | DNN (LR, SVM, PCA) | MUT, EXP | ✓ | | 10% leave-out | $r = \sim 0.86$ |
| 2019.02.27 | Li et al. [16] | Mixture (GLM, RF) | EXP | | ✓ | 20% leave-out | $r = 0.882$ |
| 2019.07.11 | Lind et al. [26] | RF (SVM, ANN) | MUT | ✓ | | 5-fold CV | $r = 0.86$ |
| 2019.07.29 | Liu et al. [34] | CNN (ANN) | MUT, CNV | ✓ | | 10% leave-out | $R^2 = 0.826$ |
| 2019.10.16 | Manica et al. [35] | CNN + RNN (RF, SVM) | EXP, CNV, PPI | ✓ | | 5-fold CV | $R^2 = 0.86$ |
| 2019.11.04 | Oskooei et al. [24] | Network (RF, LR) | EXP, PPI | ✓ | | 30-fold CV | $r = \sim 0.9$ |

ANN: artificial neural network; AUC: area under the ROC curve; CCLE: Cancer Cell Line Encyclopedia; CLS: cancer classification; CNN: convolutional neural network; CNV: copy number variation; CV: cross-validation; EN: elastic net; EXP: gene expression; GDSC: Genomics of Drug Sensitivity in Cancer; GLM: generalized linear model, including ridge, elastic net, and lasso regression; DNN: deep neural networks; LOOCV: leave-one-out cross-validation; LR: linear regression; MCC: Matthews correlation coefficient; MET: methylation; MSE: mean squared error; MUT: gene-level mutation (i.e. whether the gene is mutated or not); PCA: principal component analysis; PLS: partial least squares; PPI: protein-protein interaction; PWY: pathway; $r$ : Pearson correlation coefficient; $R^2$ : coefficient of determination; RF: random forests; $\rho$ : Spearman’s rank correlation coefficient; RNN: recurrent neural network; SGL: sparse group lasso; SNP: single nucleotide polymorphism; SVM: support vector machine.

Over the years, new techniques continue to emerge and the samples of drug response have increased constantly; nevertheless, existing prediction models still cannot achieve high performance to realize “precision” medicine goals. The prediction performances measured by the coefficient of determination (R^2^) are in the range from 0.25 to 0.78. More recently, deep neural networks with multiple hidden layers such as CDRscan [33], tCNNS [34], and MCA [35] have been proposed that achieve R^2^ higher than 0.8 (R^2^ = 0.84, 0.83, and 0.86, respectively). However, most of the cancer cell line features used in previous studies were based on gene expression profiles and did not explicitly consider associations between drugs and the structural location of mutations (Table 1). Consequently, the molecular mechanisms of drug-protein interactions cannot be inferred from these models, and thus these models have limited explainability.

The trade-off between prediction performance and explainability is an issue not only for CDRscan, tCNNS, and MCA but also for other existing machine learning approaches, thus the Defense Advanced Research Projects Agency (DARPA) recently launched the Explainable Artificial Intelligence (XAI) program [38] to facilitate building explainable models while maintaining prediction performance. In recognition of the interest in building explainable AI models, we built the Quantitative Structure-Mutation-Activity Relationship Tests (QSMART) model by (1) introducing interaction terms that capture drug-mutation relationships into quantitative structure-activity relationship (QSAR) model and using traditional statistical tests to identify significant interaction terms, and (2) utilizing a feature selection method to obtain highly informative interactions (Fig 1). Note that these interactions are statistical terms that cannot be interpreted as physical interactions directly. These two steps are equivalent to moving two hidden layers outside the neural networks “black box” for increasing the explainability of the prediction model. When applied on a subset of protein kinase inhibitors (PKIs), the QSMART achieves prediction accuracy comparable or better than state-of-the-art deep neural network methods (overall R^2^ = 0.861, AUC = 0.981, and RMSE = 0.818). Although building fully explainable models is beyond the scope of this study, we show that some level of statistically significant relationships between drugs and mutated residues contributes to drug sensitivity in cancer cell lines.

**Fig 1.**
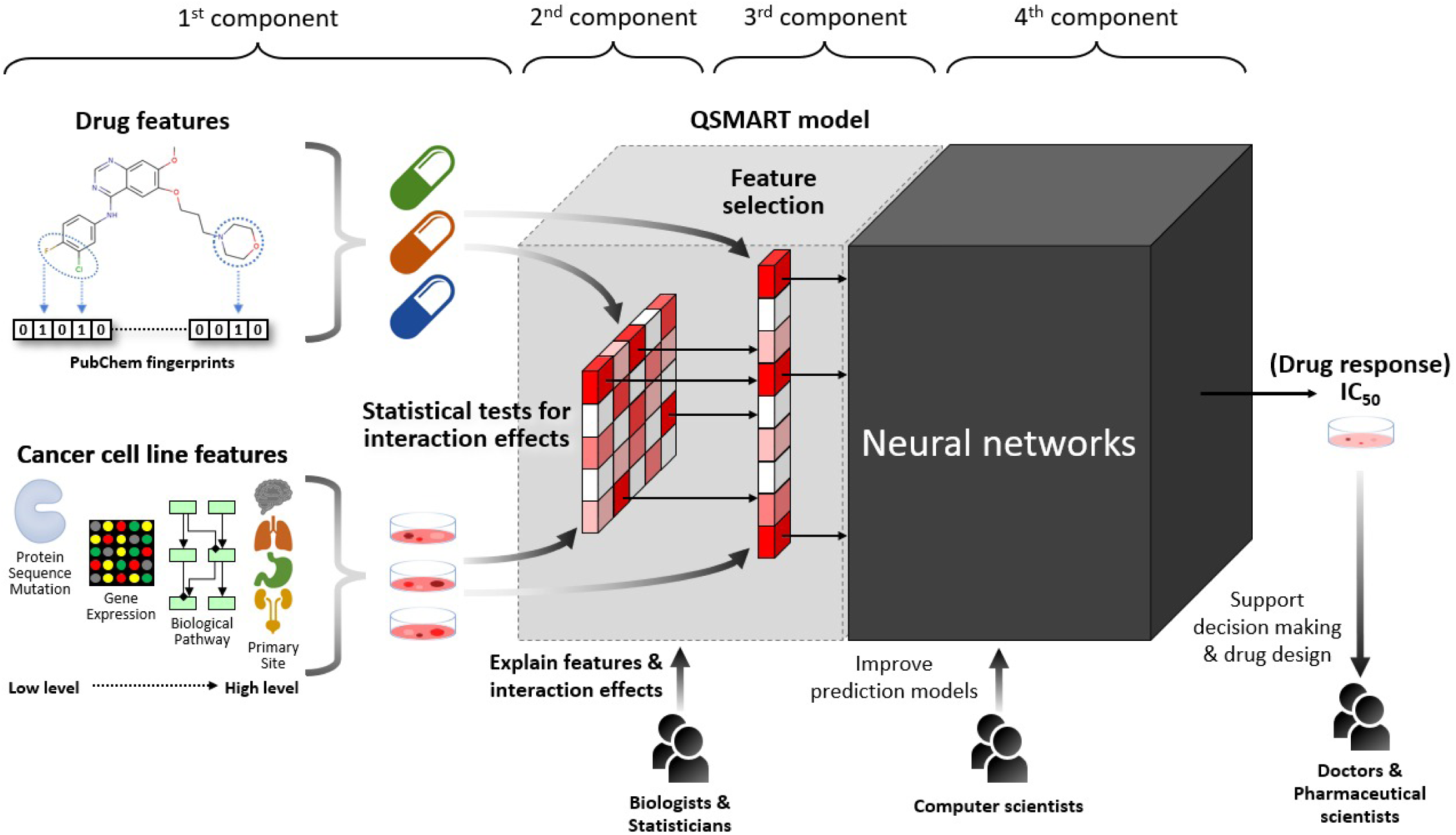
The framework of using the QSMART model with neural networks to predict protein kinase inhibitor response in cancer cell lines. Four main components of this framework: (1) drug features, cancer cell line features, and drug responses, (2) statistics tests for interaction terms, (3) a feature selection method for identifying highly informative features, and (4) a machine learning method for predicting drug response.

## Results

### Framework for protein kinase inhibitor response prediction

The overall objective of this study is to emphasize the contribution of adding interaction terms that capture drug-mutation relationships and to show how these interaction terms could help explain the mechanism of drug resistance/sensitivity. The framework we proposed in this study includes four main components: (1) the substructure fingerprints of protein kinase inhibitor (PKI) and cancer cell line’s multi-omics data, including from low-level features, such as residue mutations, to high-level features, such as perturbed biological processes, (2) F-test for identifying significant drug-mutation relationships and other interaction effects, (3) a feature selection method: Lasso with Bayesian information criterion (BIC) control, and (4) a machine learning method to predict PKI response: neural networks (Fig 1). This framework has flexibility in adapting different materials and methods in each component. To implement this framework, we collected a dataset containing ∼0.2 million drug responses (IC_50_ in a logarithmic scale; “IC_50_” hereinafter) from GDSC, split them into 23 sub-datasets according to the primary site where the cancer cell line originated, and then built a cancer-centric model for each sub-dataset. The overall prediction performance of our proposed framework and the evaluation of each component are described below.

### Overall performance of QSMART model with neural networks

For each cancer-centric model, Table 2 summarizes the number of PKI responses, the total number of features (including drug features, cancer cell line features, and interaction terms), the number of nodes in the first and second hidden layers of neural networks, and prediction performance (R^2^). Additional measurements of prediction performance (RMSE and AUC), cancer cell line features at seven feature levels, interaction terms, and training iterations are shown in S1 Table. By using the QSMART model with neural networks, we can predict the PKI response in 23 cancer types with accuracies ranging from R^2^ = 0.805 to 0.880. Fig 2a presents IC_50_ vs. predicted IC_50_ plot for all types of cancer cell lines (overall RMSE = 0.818 and R^2^ = 0.861, which means these prediction models can explain 86% of the variation in PKI responses). Although we designed three types of neural network architectures in this study: single-layer, double-layer, and complex-double-layer (see Materials and methods), we found that the prediction models for all the 23 cancer types can achieve R^2^ > 0.8 by using either single-layer or double-layer architecture. Based on Occam’s razor principle [39], we chose the simplest single-layer architecture.

**Table 2.**
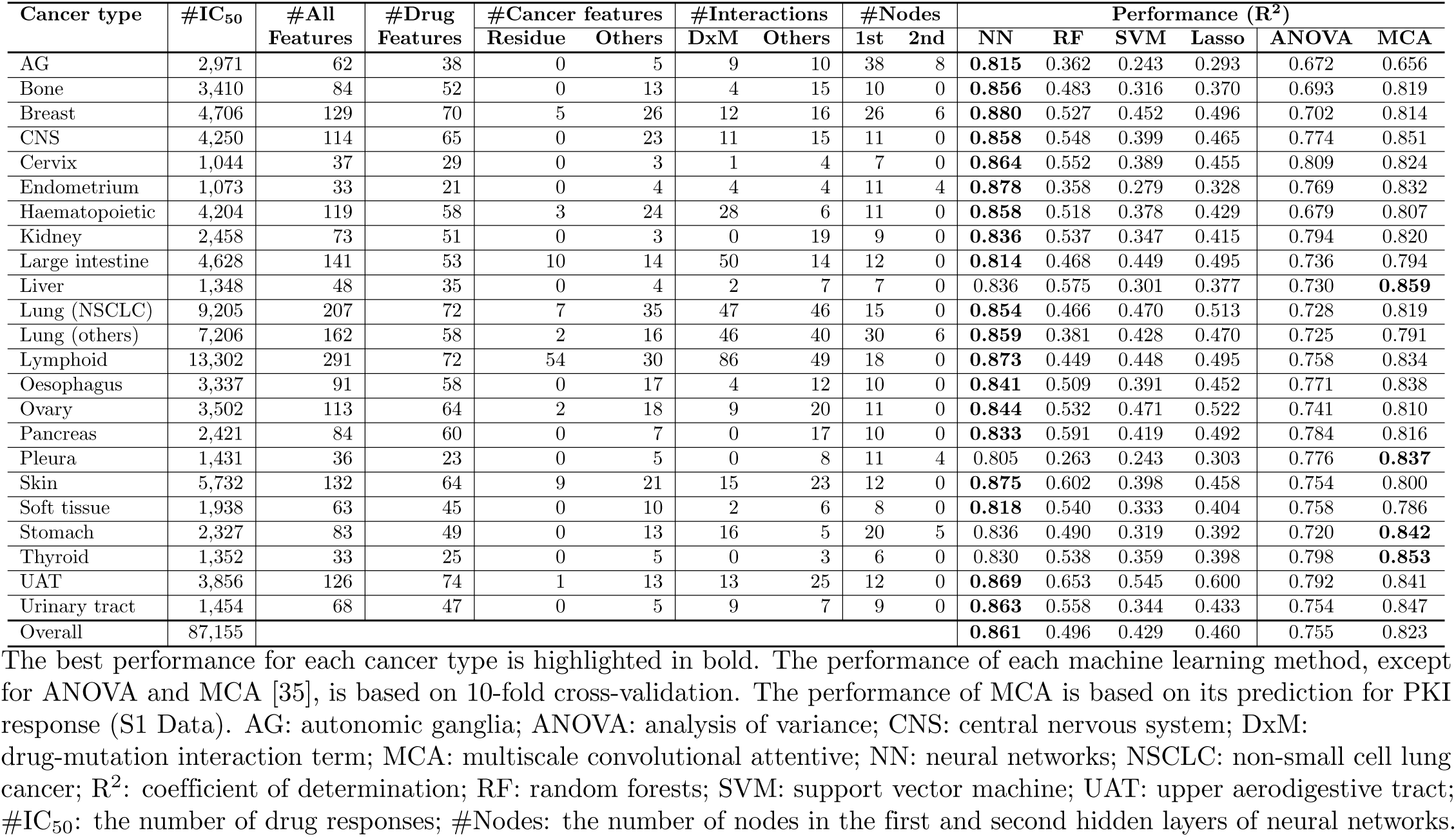
Prediction performances of using QSMART model with different machine learning methods.

**Fig 2.**
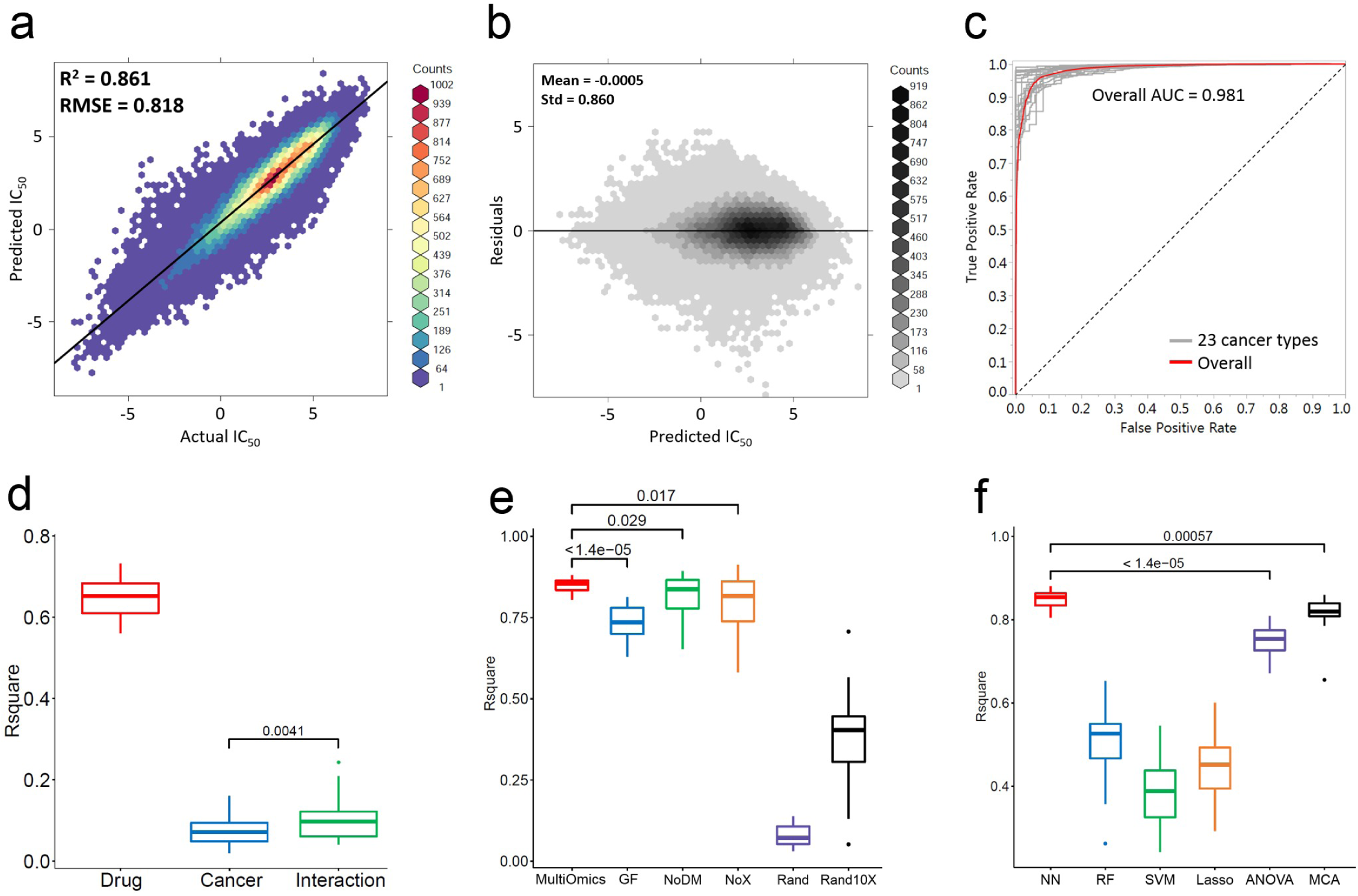
Prediction performances of different datasets and different prediction models. Wilcoxon signed-rank test is performed to compare prediction performances and the p-value is shown in each box plot. (a) Comparison between actual IC_50_ (x-axis) and the IC_50_ predicted by using QSMART with neural networks across all cancer types (y-axis). The hexbin plot is drawn by an R package “ggpubr” [40]. A fitted regression line is shown. (b) Residual analysis for the models using QSMART with neural networks across all cancer types. X-axis: predicted IC_50_; y-axis: residuals, defined as actual IC_50_ minus predicted IC_50_. (c) AUC curves of 23 cancer-centric models and an overall AUC. (d) The prediction performances of split QSMART models. (e) The prediction performances of using different datasets and different feature selection methods. GF: genomics fingerprints; NoDM: no drug-mutation interaction terms; NoX: no interaction terms; Rand: random selection; Rand10X: randomly select 10 times the number of features in the QSMART model. (f) The prediction performances of using different statistical or machine learning methods. NN: neural networks; RF: random forests; SVM: support vector machine; ANOVA: analysis of variance; MCA: multiscale convolutional attentive.

Residual analysis was then performed to assess the appropriateness of our prediction models. The overall residual plot (Fig 2b) shows that there is no specific U shape, inverted U shape, or funnel shape, which means our prediction models need no more higher-order features to capture the variation of drug responses (S1 Fig shows residual plots for all the 23 cancer types). To further confirm the prediction model’s ability to classify drug responses into two categories (sensitive vs. non-sensitive), we chose thresholds to define actual IC_50_ as sensitive or non-sensitive. Comparing to the single threshold used in a previous study [33] (IC_50_ = -2), we set multiple thresholds (−4, -3, -2, -1, and 0) and averaged the results to avoid overestimating the prediction performance. The resulting ROC curves for 23 cancer types and the overall curve are shown in Fig 2c. The overall AUC is 0.981 and comparable to a recent deep neural network-based study [33] (AUC > 0.98). AUC for each cancer-centric model is available in S1 Table.

#### Contribution of different feature categories

To roughly estimate the contribution of different feature types to the prediction accuracy, we split the features into three different categories: drug features, cancer cell line features, and interaction terms. We used the same neural network architecture (the number of nodes in the first and second hidden layers) in each cancer-centric model, and then built prediction models by using the split feature sets. Across the 23 cancer types, this experiment showed that using drug features alone to predict PKI response outperformed using cancer cell line features or interaction terms alone (overall R^2^ = 0.661, 0.126, and 0.152, respectively; Fig 2d). The contribution of interaction terms to prediction performance was significant (p-value = 0.0041, Wilcoxon signed-rank test) in comparison to cancer cell line features. Although it was partially due to the number of selected drug features being more than those of the other two feature categories, the main reason was that the drug features were more informative in cancer-centric models. Since the entire training dataset was split into 23 cancer-centric datasets, the similarity among cancer cell lines in one dataset was higher than the similarity among PKIs and thus the drug features had higher variation and higher entropy.

Assuming that the features from different categories in a full model are independent and can explain the variation of drug response independently, the summation of the prediction performances of split models (the R^2^_SSP_ in Table 3) would ideally be the upper limit of a full model. However, Table 3 shows that the prediction performances R^2^_Full_ are even higher than R^2^_SSP_ for 14 cancer types, which implies that the synergistic prediction performances (R^2^_Full_ - R^2^_SSP_) are potentially derived from the higher-order interactions performed by neural networks. Interestingly, we found that the neural network architectures of the models with the top four synergistic effects are all double-layer neural networks, instead of single-layer neural networks, which also supports our hypothesis that the synergistic prediction performance is derived from higher-order interactions.

**Table 3.**
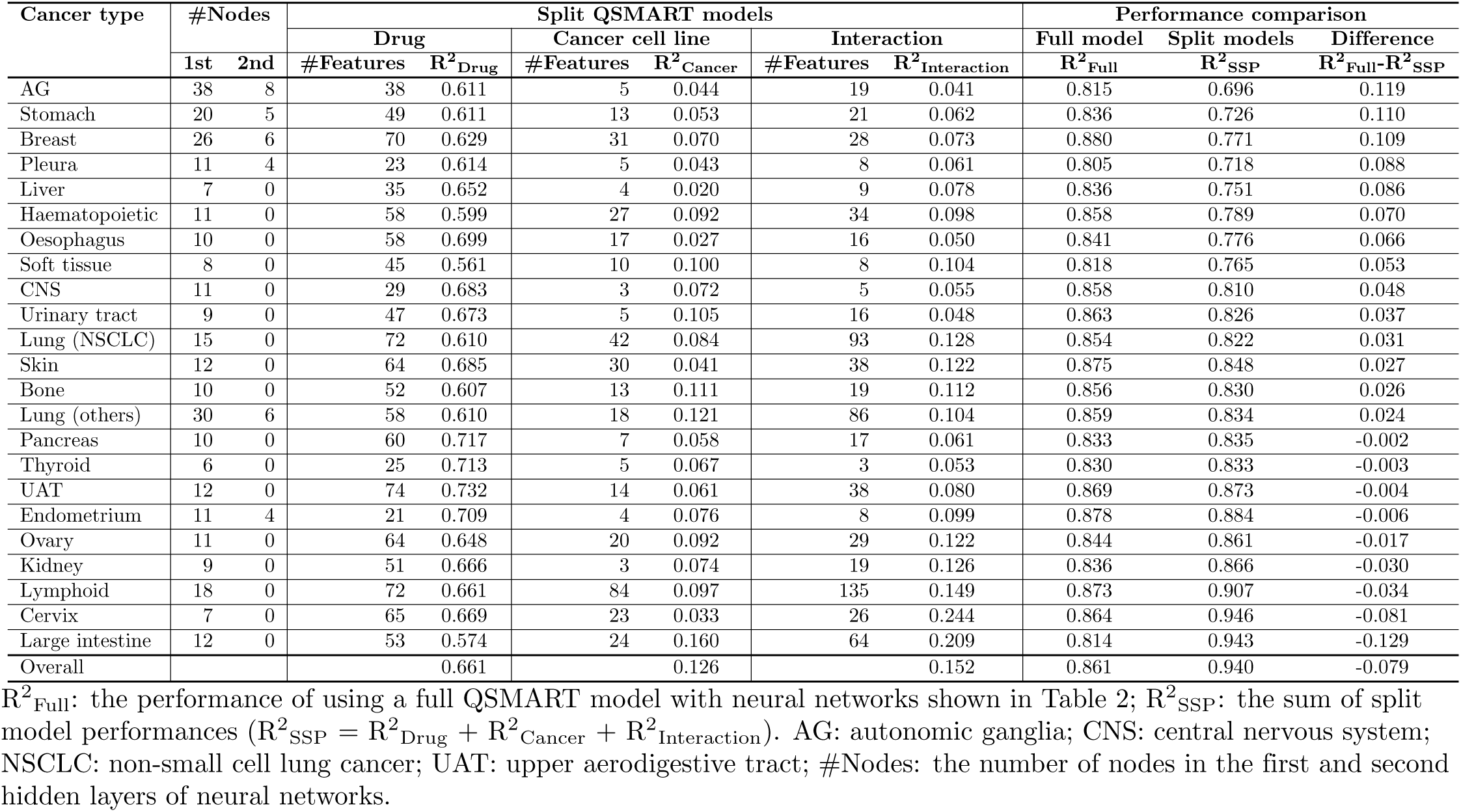
Prediction performances of using split QSMART models with neural networks.

### Informative features for predicting PKI response: multi-omics data

To evaluate the first component of the framework in this study – drug features and cancer cell line’s multi-omics data – referring to the features used in a previous study [33] which achieves a high prediction accuracy, we replaced our drug features and cancer cell line features with 3,072 molecular fingerprints and 44,364 genomic fingerprints (S2 Fig). According to the previous study’s method, drug fingerprints were generated by PaDEL-descriptor [41] (a software to calculate molecular descriptors and fingerprints) and genomic fingerprints (genomic mutation positions) were obtained from COSMIC Cell Lines Project [42]. To make them comparable with our models, we used the same feature selection method to prioritize all the drug fingerprints and genomic fingerprints, selected the same total number of features in our model for each cancer type (shown in Table 2), and then used the same neural network architectures. The number of selected features and prediction performances are shown in S2 Table. The box plot in Fig 2e shows that the performance distribution of 23 cancer-centric models using multi-omics data is significantly higher than that of the models using genomic fingerprints (overall R^2^ = 0.861 vs. 0.752, p-value < 1.4e-05, Wilcoxon signed-rank test). Under the same sample size, the number of features, and neural network architectures, our experiment shows that compared to genomic fingerprints, multi-omics data are more informative features for predicting PKI response.

### Contribution of interaction terms in the QSMART model

To evaluate the second component of the framework in this study – F-test for identifying significant interaction terms – we performed two experiments: removing drug-mutation interaction terms and removing all interaction terms. We utilized the feature selection method to prioritize all input features, selected the same total number of features in the original models shown in Table 2, and then used the same neural network architectures to train the new models. The results of these two experiments are shown in S3 Table and S4 Table, respectively. The box plot in Fig 2e shows that the full QSMART models significantly outperform the models without drug-mutation interaction terms (overall R^2^ = 0.861 vs. 0.839, p-value = 0.029) and the models without any interaction terms (overall R^2^ = 0.861 vs. 0.823, p-value = 0.017). Interestingly, compared to the full QSMART models, we found that the prediction models of some cancer types, such as upper aerodigestive tract and breast, achieved higher performance without using any interaction terms. We conjectured that it was because some informative high-order interactions were captured inside the neural network black box and thus compensated the lack of interaction terms in the input layer. However, using neural networks cannot guarantee that these informative but unexplainable high-order interactions will be captured under the limited number of samples and the training iteration we used. This fact is reflected in Fig 2e, which shows that the prediction performance is variable when the drug-mutation interaction terms are not used (R^2^ = 0.653 to 0.892) or all the interaction terms are not used (R^2^ = 0.581 to 0.912). In this paragraph, we validate that adding interaction terms significantly contributes to PKI response prediction; in a non-small cell lung cancer case study described below, we will illustrate how the statistical interaction terms are interpreted as potential physical drug-target interactions and thus increase the explainability of prediction models.

### Feature selection method: Lasso with BIC control

We next evaluated the feature selection methods. Since building neural networks using all the features from the first two components of the framework cannot be finished within a reasonable amount of time, instead of removing the third component of the framework, we randomly selected the same number of features in the original model for each cancer type shown in Table 2. Then, we trained a new neural network model with the same architecture for each cancer type. For each cancer type, the number of randomly selected features along with prediction performances are shown in S5 Table. It is not surprising that the prediction performances dropped to R^2^ = 0.031 to 0.138 (overall R^2^ = 0.125). To further evaluate the feature selection method we used, we increased the number of randomly selected features to 10 times the original number. The performances increased to R^2^ = 0.052 to 0.707 (S6 Table; overall R^2^ = 0.378). Nevertheless, these performances are significantly lower than those of the original models that utilize Lasso with BIC control (Fig 2e). Although the second component of the framework in this study had provided a lot of statistically significant interaction terms, compared to random selections, Lasso with BIC control identified highly informative features for PKI response prediction.

### Comparison of machine learning methods used in our framework

We next compared neural networks with three commonly used machine learning models for drug response prediction: random forests, SVM, and Lasso regression. Based on the same feature set as inputs, neural networks significantly outperformed other machine learning approaches (Table 2 and Fig 2f; overall R^2^ = 0.861, 0.496, 0.429, and 0.460 for neural networks, random forests, SVM, and Lasso regression, respectively).

Furthermore, based on the feature sets used to validate the contribution of previous components in the framework, neural networks also outperformed random forests, SVM, and Lasso regression (S2 Table-S6 Table). A previous study [33] also showed that neural networks outperformed random forests and SVM (R^2^ = 0.843, 0.698, and 0.562 for DNN, random forests, and SVM, respectively) in drug response prediction. Interestingly, neural network was only slightly better than Lasso in overall performance when randomly selected features were used as inputs (R^2^ = 0.125 vs. 0.116, p-value = 0.015, Wilcoxon signed-rank test).

In addition to the three commonly used machine learning approaches, we compared our models with two-way ANOVA analysis and MCA [35]. Two-way ANOVA analysis was applied to assess the ability of two factors – drug and cancer cell line – to explain the variation of drug response. Drug IDs and cancer cell line IDs represented different levels of drug and cancer cell line, respectively. The result of two-way ANOVA showed that these two factors could explain 67.2% to 80.9% of the drug response variation in different cancer types (Table 2; overall R^2^ = 0.755). Compared to ANOVA and MCA, using the QSMART model with neural networks had significantly higher ability to explain the PKI response variation for 23 cancer types (p-value < 1.4e-05 and p-value = 0.00057, respectively; Fig 2f).

### Case study: non-small cell lung cancer

We next evaluated the contribution of different features in drug response prediction using non-small cell lung cancer as a case study. All 207 features in the NSCLC-specific QSMART model and their descriptions are listed in S2 Data. We choose several pertinent features and explain their biological relevance in this case study to demonstrate how scientists may use our prediction model to explain their findings.

#### Drug feature

“From Sanger”. This feature was introduced into the model to distinguish the assays done by Massachusetts General Hospital (0) or Wellcome Sanger Institute (1). This feature represents the batch effects among the laboratory experiments performed by these two institutes. On average, the PKI responses obtained from Massachusetts General Hospital showed lower drug sensitivity (higher IC_50_ value) than those from the Wellcome Sanger Institute in the NSCLC dataset (average actual IC_50_ = 2.88 vs. 2.41). To investigate these experimental batch effects, we increased one unit to this feature and held other features constant. Although holding other features constant is not possible in reality, from the mathematical point of view, the result showed that if we replace 0 with 1 for From Sanger, the average IC_50_ predicted by our pre-trained model will reduce 0.65 (average predicted IC_50_ = 2.87 vs. 2.22, S2 Data). Interestingly, this feature was selected not only in the NSCLC model but also in other 22 cancer-centric models, meaning the batch effects were significant across the assays done by these two institutes.

#### Biological processes interaction

“GO 0030324 X GO 0048675”. This feature represents the multiplication of the number of mutations perturbing the biological process “lung development” (Gene Ontology ID: GO:0030324) and the number of mutations perturbing “axon extension” (Gene Ontology ID: GO:0048675). Axon initiation, extension, and guidance are known to play important roles in cancer invasion and metastasis [43]. In the NSCLC dataset, there are eight cell lines with mutations in protein kinases associated with axon extension; among them, NCI-H1944 and NCI-H2030 are from patients with metastatic NSCLC. On average, the NSCLC cell lines with this interaction showed higher PKI responses than those without this interaction (average actual IC_50_ = 4.32 vs. 2.69) and those involved in “lung development” or “axon extension” alone (average actual IC_50_ = 3.20 or 2.07, respectively). Based on our prediction model, every unit increase in this interaction term is associated with a 0.45 unit increase in IC_50_ on average (average predicted IC_50_= 2.73 vs. 3.18).

#### Protein-protein interaction

The NSCLC model contains 27 protein-protein interaction (PPI) terms. Each PPI was quantified by the product of the participants’ gene expression level. Every unit of gene expression level increase in these 27 PPIs is contributed to -0.089 to 0.061 unit increase in IC_50_ on average. Gene enrichment analysis of the 27 genes in the TP53-centric subnetwork shown in Fig 3 revealed the overrepresentation of pathways associated with angiogenesis, inflammation, apoptosis, and axon guidance (performed by PANTHER [44], S7 Table). MAP4K4, one of the genes involved in the apoptosis signaling pathway, is an emerging therapeutic target in cancer [45], and its over-expression is a prognostic factor for lung adenocarcinoma [46]. MAP4K4 expression is up-regulated upon binding by p53, and it will then activate the JNK signaling pathway to drive apoptosis [47]. In the NSCLC dataset, when the expression of MAP4K4-TP53 interaction increases, average IC_50_ is slightly decreased (Pearson correlation = -0.10); in our pre-trained PKI response prediction model, every unit of gene expression level increase in MAP4K4-TP53 PPI is associated with 0.012 unit decrease in IC_50_ on average (average predicted IC_50_ = 2.727 vs. 2.715).

**Fig 3.**
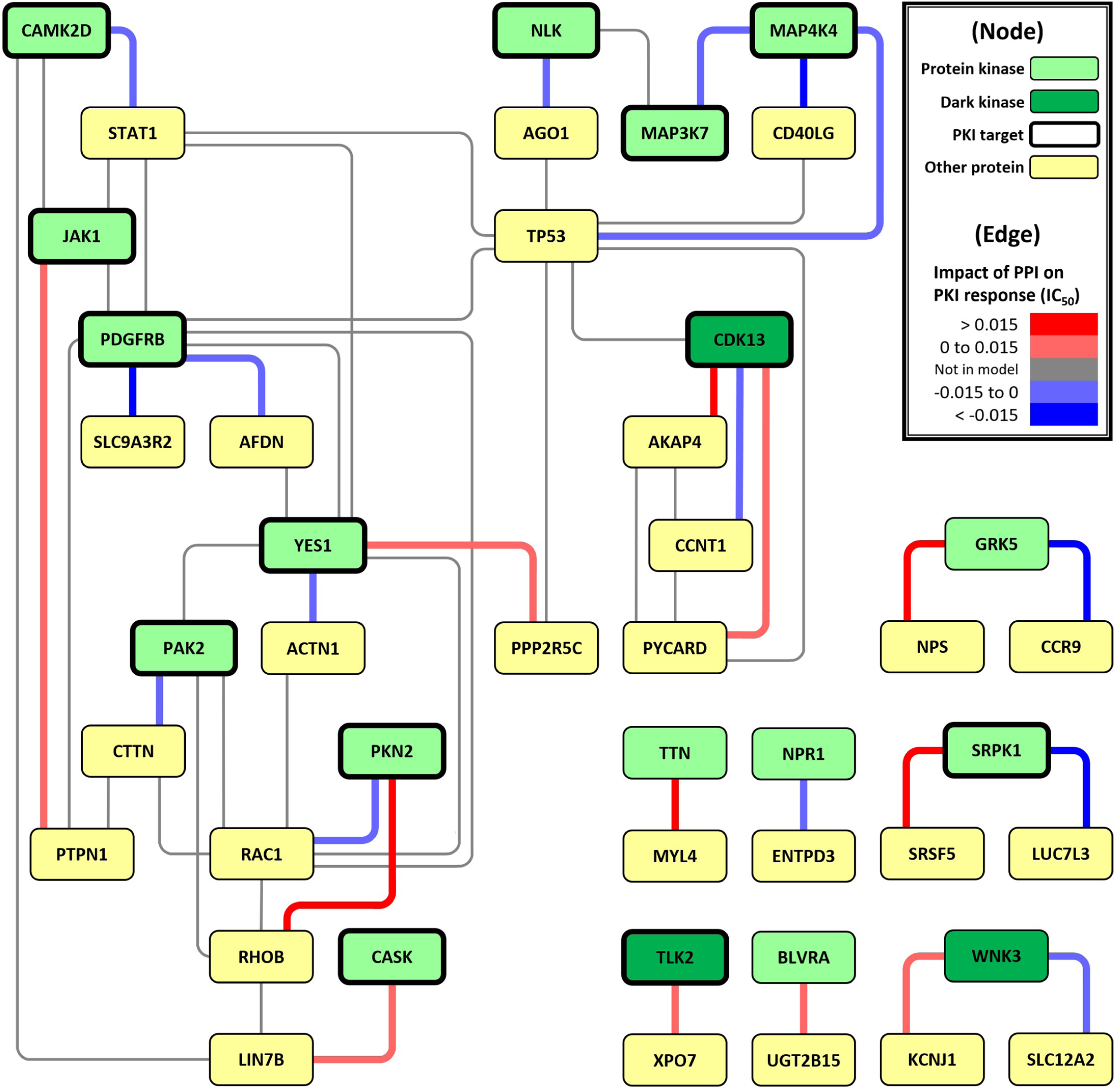
Protein-protein interaction network constructed by the interaction terms for predicting PKI response in NSCLC cell lines. Green node: protein kinase; dark green node: dark/understudied protein kinase; yellow node: other protein; the node with a thick border: known PKI target; red edge: the PPI with a positive impact on IC_50_; light red edge: the PPI with a weak positive impact on IC_50_; blue edge: the PPI with a negative impact on IC_50_; light blue edge: the PPI with a weak negative impact on IC_50_; gray edge: the PPI not in the prediction model.

CDK13, an understudied protein kinase defined by NIH Illuminating the Druggable Genome (IDG) program [48] (S3 Data, last updated on June 11, 2019), participates in a 4-clique PPI module in the TP53-centric subnetwork (Fig 3). Its three PPIs in this module are all the features of the NSCLC-specific model. One of CDK13’s PPI partners, AKAP4, is a biomarker for NSCLC, and its expression increase was associated with tumor stage [49]. In addition to NSCLC, AKAP4 is also a potential therapeutic target of colorectal cancer [50] and ovarian cancer [51], and it regulates the expression of the CDK family. In the NSCLC dataset, the expression of CDK13-AKAP4 interaction had a weak positive correlation with IC_50_ (Pearson correlation = 0.07); in the prediction model, every unit of gene expression level increase in CDK13-AKAP4 PPI is associated with 0.017 unit increase in IC_50_ on average (average predicted IC_50_ = 2.727 vs. 2.744).

#### Drug-mutation interaction

In total, there are 47 drug-mutation interaction terms in the NSCLC model, and they are located at 22 PKA positions represented by spheres in Fig 4a (PDB ID: 1ATP). Their impacts on IC_50_ are listed in S8 Table, sorted by absolute IC_50_ impact. The drug-mutation relationships located in the canonical ATP-binding pocket (highlighted by a dashed rectangle in Fig 4a) could be formed by type I or type II protein kinase inhibitors according to the protein structure’s active or inactive conformation, respectively [52].

**Fig 4.**
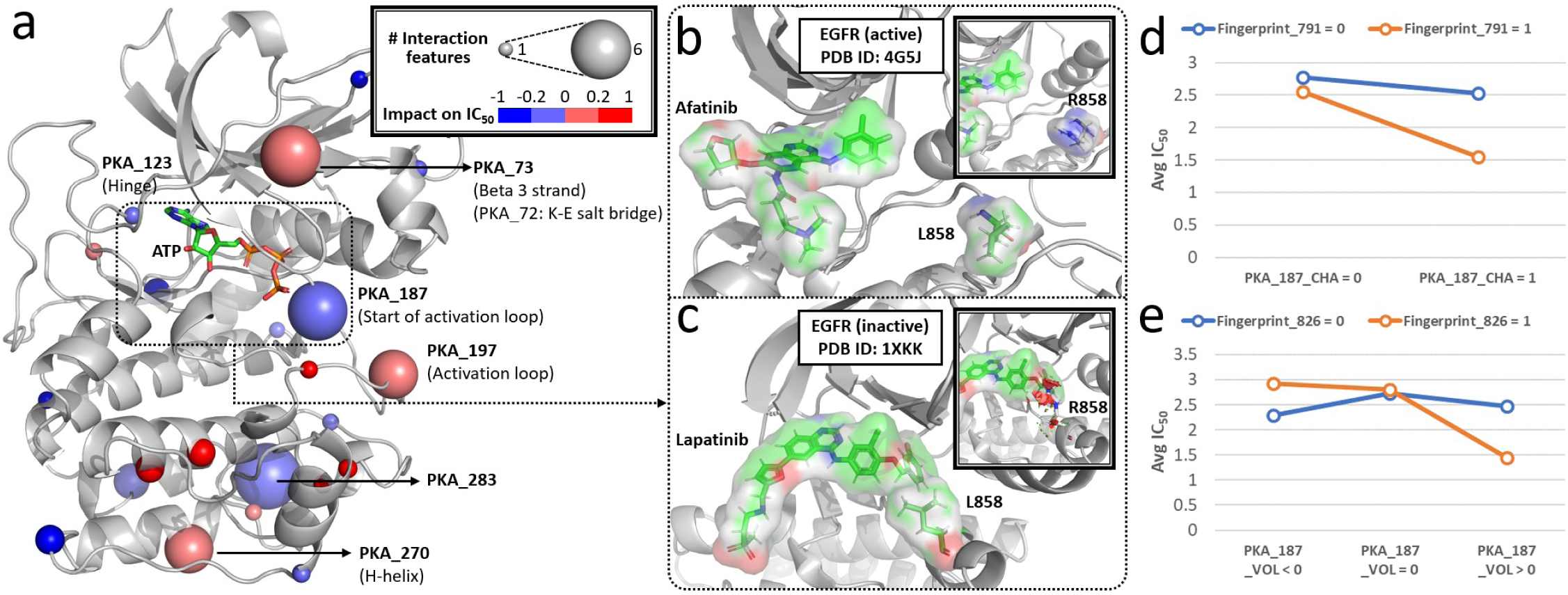
Distribution of drug-mutation relationships on the reference protein kinase A structure and interaction analyses. (a) Interaction hot spots are labeled and represented by larger spheres on the reference PKA structure (PDB ID: 1ATP, visualized by PyMol [53]). If a residue involved in multiple drug-mutation relationships, the median of their impacts on IC_50_ is chosen to represent the color of the sphere. Red sphere: the drug-mutation relationship with a positive impact on IC_50_; light red sphere: the relationship with a weak positive impact on IC_50_; blue sphere: the relationship with a negative impact on IC_50_; light blue sphere: the relationship with a weak negative impact on IC_50_. (b) and (c): Examples of two PKIs (afatinib and lapatinib) with different binding modes in the active (PDB ID: 4G5J) and inactive (PDB ID: 1XKK) conformations of EGFR crystal structures, respectively. The residue corresponding to PKA_187 – EGFR L858 – is labeled in each example; its arginine mutant form simulated by PyMol is illustrated. (d) and (e): Statistical interaction analyses for Fingerprint_791 vs. PKA_187_CHA and Fingerprint_826 vs. PKA_187_VOL in the NSCLC dataset, respectively.

Fig 4b and Fig 4c respectively illustrate different binding modes of two EGFR inhibitors in our dataset (afatinib and lapatinib) that contribute to variable response in L858R mutant EGFR (L858 corresponds to position 187 in protein kinase A; PKA_187).

H3255, an NSCLC cell line with EGFR L858R mutation, is hypersensitive to afatinib treatment (IC_50_ = -4.35; average IC_50_ = 2.03 for all the NSCLC cell lines treated with afatinib). Notably, the L858R mutation can be accomodated in the active conformation of EGFR, but not in the inactive state due to steric hindrance [54].

An interaction analysis (Fig 4d) shows that the mutated residues involving charge difference at PKA_187 have significant interaction (p-value = 0.043, F-test) with Fingerprint 791, a drug substructure “NC1CCC(N)CC1” of afatinib). Based on our prediction model, every unit increase in the interaction term “PKA_187_CHA_X_Fingerprint 791”, an interaction term with one of the highest impact on IC_50_ among all the drug-mutation interaction terms in the model (S8 Table), is associated with a 0.46 unit decrease in IC_50_ on average (average predicted IC_50_ = 2.73 vs. 2.27). Another interaction analysis (Fig 4e) shows that the mutated residues involving volume difference at PKA_187 have significant interaction (p-value = 0.035, F-test) with Fingerprint 826 (a drug substructure “OC1C(N)CCCC1” of afatinib). Every unit increase in “PKA_187_VOL_X Fingerprint_826” is associated with a 0.01 unit decrease in IC_50_ on average (average predicted IC_50_ = 2.73 vs. 2.72). Since lapatinib lacks both substructures Fingerprint_791 and Fingerprint_826, we speculate that mutant EGFR in NSCLC cells with a larger, positively charged mutation at PKA_187 are resistant to lapatinib (the blue lines in Fig 4d and Fig 4e).

## Discussion

In this study, we propose a PKI response prediction framework to accurately estimate IC_50_ values with a more explainable AI model. This framework includes four components: (1) drug features, cancer cell line’s multi-omics data, and PKI responses, (2) statistical tests for capturing interaction effects, (3) feature selection, and (4) neural networks. We validate the contribution of each component, showed high prediction performances, and used NSCLC dataset as a case study to explain several features. We systematically investigate the contributions of various interaction effects (such as protein-protein interactions, pathway-pathway interactions, and drug-mutation relationships) to drug response prediction.

The intrinsic limitation of drug response prediction is the unexplainable variation of drug response caused by different assays and experimental conditions. Currently, GDSC and CCLE are the two main sources for studying cancer drug response. Several previous studies about predicting drug response used data not only from GDSC but also from CCLE (Table 1). However, a previous study [20] pointed out that although GDSC and CCLE datasets shared 343 cancer cell lines and 15 drugs, the drug responses from these two datasets were poorly correlated. Thus, we chose to only use a single source in this study to minimize the unexplainable effect from different experimental environments. Nevertheless, this situation impeded us from finding appropriate independent testing set outside the GDSC data. Even though the drug response data we used were only from GDSC, the feature selection process showed that the drug feature “From Sanger” was selected for all the 23 cancer-centric prediction models, meaning the batch effects were significant across the assays. While our studies were underway, we noticed that GDSC 8.0 was released. Compared with release 7.0, it contains 160 thousand more drug responses. However, this dramatic increase did not provide us a syncretic testing set since the old drug response dataset (called GDSC1 in release 8.0) and the new drug response dataset (called GDSC2) were generated based on different types of assays. Although the drug responses measured by different assays seem to have high correlation (R = 0.838 in Pearson correlation coefficient), unfortunately, it implies that even if we train a perfect model for GDSC1, the performance of predicting the drug responses in GDSC2 as an independent testing set would only be R^2^ = 0.702 (S3 Fig panel a). Moreover, if we only focus on PKI responses between the two datasets, the correlation is reduced to 0.774 and R^2^ = 0.599 (S3 Fig panel b). Furthermore, if we use our pre-trained models to predict the PKI response in GDSC2, the overall performance drops to R^2^ = 0.556 (S3 Fig panel c).

In the case study, we illustrated the possibility of interpreting statistical interaction terms into potential physical interactions. When we investigated the contribution of protein-protein interactions to drug response prediction, the original purpose of utilizing biological knowledge (such as known PPIs from STRING [55]) was to narrow down the huge search space (a matrix of 30,000 proteins by 30,000 proteins). Consequently, this additional information also helped us explain the biological role of these statistical interaction terms with preliminary evidence. On the contrary, when we investigated the contribution of drug-mutation relationships to drug response prediction, we explored all the interactions between drug features and the mutations at reference PKA positions. Although restricting the mutations to be in the region around ATP-binding pocket (from PKA_47 to PKA_188, defined by the Kinase-Ligand Interaction Fingerprints and Structures (KLIFS) database [56]) could increase the probability of finding physical interactions among those statistical interaction terms, we will lose the opportunity to explore the potential interactions between PKIs and the allosteric binding sites outside the ATP-binding pocket.

In conclusion, by integrating multi-omics data, utilizing the innovative QSMART model, and employing neural networks, we not only can accurately predict PKI responses in cancer cell lines but also increase the explainability behind our prediction models. Compared to traditional QSAR models, the QSMART model proposed in this study further introduces different types of interaction terms. While we demonstrate our model in protein kinase binding, the QSMART model can be applied to other protein families, such as G protein-coupled receptors (GPCRs) and ion channels. Moreover, the concept of QSMART model can also be broadly applied to other types of interactions, such as the protein-protein interaction that we had demonstrated, drug-drug interaction, glycosyltransferase-donor analog interaction, gene-environment interaction, and agent-host interaction.

## Materials and methods

### Protein kinase inhibitor

We define small-molecule (molecular weight < 900 daltons) protein kinase inhibitors (PKIs) in GDSC (release 7.0) [8] from a variety of publicly available, manually curated drug-target databases, and experimental data. The list of human protein kinases in this study is defined by ProKinO (version 2.0) [57]. Drug-kinase associations were extracted from DrugBank (version 5.1.0) [58], Therapeutic Target Database (TTD, last accessed on September 15th, 2017) [59], Pharos (last accessed on May 15th, 2018) [60], and LINCS Data Portal (last accessed on May 15th, 2018) [61]. We define a drug as a PKI if it is annotated as an “inhibitor”, “antagonist”, or “suppressor” in the drug-kinase associations. We also include the PKIs in LINCS Data Portal if their controls are less than 5% in KINOMEscan^®^ assays. Based on these criteria, we define 143 small-molecule PKIs out of the 252 unique screened compounds in GDSC (S4 Data).

### Drug response

GDSC provides the half-maximal inhibitory concentration values (IC_50_, on a logarithmic scale) for 224,202 drug-cancer cell line pairs of drug sensitivity assays. These assays were performed by either the Wellcome Trust Sanger Institute or Massachusetts General Hospital Cancer Center. In this drug response dataset, there are 12,509 duplicated drug-cancer cell line pairs derived from 16 duplicated drugs. We measured the Pearson correlation coefficient between the IC_50_ values of each duplicated drug. Only afatinib and refametinib showed a strong positive correlation (r > 0.7); their IC_50_ values were then merged by their weighted means [62]. Drug responses of all other duplicated drugs were excluded from our study as they may have been assayed under different experimental conditions. The resulting dataset of 197,459 non-redundant drug responses consists of 236 drugs and 1,065 cancer cell lines. After filtering out non-PKIs, 109,856 non-redundant drug responses consisting of 135 PKIs and 1,064 cancer cell lines remained.

### Drug features

Drugs’ 2D structures were obtained from PubChem in SDF format. The CDK Descriptor Calculator GUI (version 1.4.6) [63] generated 881 PubChem fingerprints and 286 chemical descriptors including constitutional, topological, electronic, geometric, and bridge descriptors. Observing high multicollinearity within features, we removed redundant features and implemented the variance inflation factor (VIF) criterion [64] to reduce multicollinearity (for more details, see the Feature screening section below). After filtering, 92 PubChem fingerprints and 0 chemical descriptors remained.

To compare our prediction performances with those in a previous study [33], we used the same methods to generate (1) fingerprints, (2) extended fingerprints, and (3) graph-only fingerprints by PaDEL-descriptor (version 2.21) [41] for each drug. In total, there are 3,072 binary descriptors as drug features in the compared training sets. The relatively large, unfiltered set of drug features are only used for comparison purposes in our study.

### Cancer cell line features

Using mutation profiles for each cancer cell line sample provided by COSMIC Cell Lines Project (v87) [42], we incorporate 7 categories of multi-omics data to quantify the differences between wild type and mutant protein kinases: (1) residue-level: reference protein kinase A (PKA) position (from ProKinO), mutant type, charge, polarity, hydrophobicity, accessible surface area, side-chain volume, energy per residue [65], and substitution score (BLOSUM62 [66]); (2) motif-level: sequence and structural motifs of protein kinase (from ProKinO); (3) domain-level: subdomain in protein kinase (from ProKinO) and functional domain (from Pfam v31.0 [67]); (4) gene-level: the number of mutations in the genes encoding protein kinases, gene expression (from GDSC), and copy number variation (from COSMIC); (5) family-level: protein kinase family and group (from ProKinO); (6) pathway-level: reaction, pathway (from Reactome [68], last accessed on May 15th, 2018), and biological process (from AmiGO [69], last accessed on May 15th, 2018); and (7) sample-level: microsatellite instability, average ploidy, age, cancer originated tissue type, and histological classification (from COSMIC and Cellosaurus [70]).

The formula for generating all cancer cell line features is shown in S9 Table. Residue-level features of a cancer cell line were extracted from COSMIC mutants labeled as “Substitution - Missense”. These features were then calculated if the mutation position could be aligned to the reference PKA position. This choice is based on an assumption that, for all protein kinases, mutations at equivalent positions will have similar effects on drug response. We further used two different types of weights, conservation score (KinView [71] with Jensen-Shannon divergence calculation [72]) and gene expression, to estimate the different effects of the same mutant type aligned to the same PKA position from different protein kinases.

Based on mutation position, the values of motif-level or domain-level features were calculated if it occurs in a specific motif or domain and its mutation description is “Substitution - Missense” or in-frame INDELs (insertions and deletions) in COSMIC. All mutation types, except for “Substitution - coding silent” and “Unknown”, were taken into account for calculating the values of gene-level and higher-level features. For missing data, we assigned “Neutral” for copy number variation and “Unknown” for microsatellite instability and gender. No imputation was implemented for missing age.

### QSMART model

The Quantitative Structure-Mutation-Activity Relationship Tests (QSMART) model was developed based on the QSAR model a variety of interaction terms. First, we built a basic model with all drug features and cancer cell line features as independent variables for estimating IC_50_:

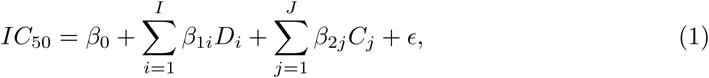

where *β*_0_ is the intercept, *β*_1*i*_ and *β*_2*j*_ represent the coefficients of the *i*th drug feature *D*_*i*_ and the *j*th cancer cell line feature *C*_*j*_, and *ϵ* is the error term.

Because the residue-level features of a cancer cell line represent the mutation status in the reference PKA structure and we are interested in investigating drug-mutation relationships, we introduced drug-mutation interaction terms into the model:

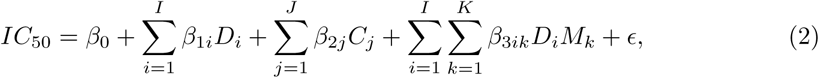

where *β*_3*ik*_ is the coefficient of the interaction term formed by the *i*th drug feature *D*_*i*_ and the *k*th residue-level feature *M*_*k*_. Since all cancer cell line features contain residue-level features and other six feature categories, {*C*_1_, …, *C*_*J*_} is a superset of {*M*_1_, …, *M*_*K*_}. Considering that the interaction terms formed by the substructures of drug and high-level cancer cell line features have no biological relevance, we did not incorporate all cancer cell line features as part of interaction terms. For example, we did not consider the interaction between a substructure “Fingerprint 1” and a biological process “lung development” because it is unexplainable.

In addition to using all cancer cell line features, we further introduced more types of interaction terms into the full QSMART model to capture the environment of a cancer cell line:

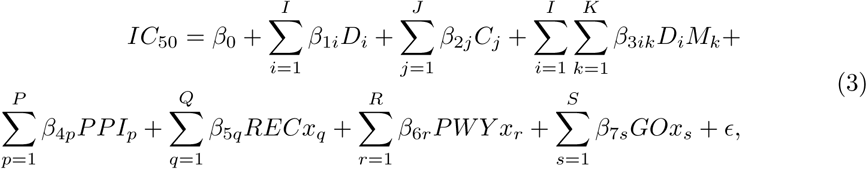

where *β*_4*p*_, *β*_5*q*_, *β*_6*r*_, and *β*_7*s*_ are the coefficients of the *p*th protein-protein interaction *PPI*_*p*_, the *q*th reaction-reaction interaction *RECx*_*q*_, the *r*th pathway-pathway interaction *PWY x*_*r*_, and the *s*th biological processes interaction *GOx*_*s*_, respectively. These four types of interaction terms are formed by all pairs of protein, reaction, pathway, and biological process features, respectively. More details about interaction terms are described below.

### Interaction terms

Five types of interaction terms were introduced into the QSMART model: drug-mutation interaction, protein-protein interaction, reaction-reaction interaction, pathway-pathway interaction, and biological processes interaction. These interactions were not necessarily physical interactions; instead, they were predictors that show statistically significant contribution to explaining the variation of IC_50_ values. For drug-mutation interaction term, only the residues mapping to the reference PKA structure were considered to form interactions with drugs. To reduce the search space, prior biological knowledge was used to filter interactions with less biological relevance. For protein-protein interaction (PPI), we retain PPIs with scores higher than 700 in the STRING database [55]. Gene expression level was used as a weight for PPIs to roughly represent the protein abundance in cancer cell lines. For reaction, pathway, and biological processes interactions, we removed the interactions formed by two entities from the same biological process/pathway hierarchy. For instance, the interaction between the biological process “lung cell differentiation” (GO:0060479) and its parent “lung development” (GO:0030324) was removed since it is unexplainable. Each interaction term was tested individually by F-test using R (version 3.4.4) [73]. Significant interaction terms (FDR < 0.05) with no less than 30 non-zero values were taken for further feature selection.

### Datasets

To reduce more potential sources of noise and bias, we further filter cancer cell lines from the PKI response dataset if (1) their mutation profiles are not detected by whole-genome sequencing,(2) they have less than 30 drug response entries, (3) their gene expression profile is not available, or (4) their mutation site does not map to a residue in the reference PKA position. The dataset was then split into 29 groups, stratified by cancer primary site. Groups with less than 1,000 responses (adrenal gland, biliary tract, placenta, prostate, salivary gland, small intestine, testis, and vulva) were excluded due to low statistical power. “Haematopoietic and lymphoid tissue”, the largest group, was further divided into two subsets by primary histology: “haematopoietic neoplasm” and “lymphoid neoplasm”. For the case study, we collected cancer cell lines for the non-small cell lung cancer (NSCLC) dataset from the lung cancer dataset if their histology subtype is adenocarcinoma, non-small cell carcinoma, squamous cell carcinoma, large cell carcinoma, giant cell carcinoma, or mixed adenosquamous carcinoma. Remaining lung cancer cell samples were classified as “lung (others)”. We created cancer type-centric training sets by expanding the drug response dataset with drug features, cancer cell lines features, and significant interaction terms. Categorical data in the training sets were coded into dummy variables. As a result, we prepared 23 cancer type-centric training sets. The number of PKI response for each cancer type is shown in Table 2.

### Feature screening

Observing high multicollinearity within the features in the first component of our prediction framework (Fig 1), we implemented the variance inflation factor (VIF) criterion [64] to remove highly correlated features. For the multiple regression model with *f* features, *X*_*i*_ (*i* = 1, …, *f*), the VIF for the *i*th feature can be expressed by:

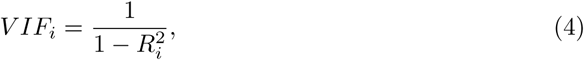

where 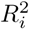 is the coefficient of determination of the regression between *X*_*i*_ and the remaining *f* − 1 features. *V IF*_*i*_ > 5 (i.e. 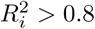) is considered to be high collinearity [74] and *X*_*i*_ should be excluded from the model. We first prioritized drug features based on these rules: (1) the later PubChem fingerprint bit positions (complex patterns) have higher priorities than the earlier ones (simple elements), and (2) PubChem fingerprints have higher priorities than calculated chemical descriptors because fingerprints directly represent molecular substructures of the drug. Then, starting from higher priority features moving towards lower priority features, we implemented stepwise selection under VIF control.

Co-expressed genes in the same prediction model also exhibited collinearity. To address this issue, we also used the VIF criterion to filter co-expressed genes in each training set. We prioritize genes based on the Pearson correlation coefficient between their expression and IC_50_ values, then implemented stepwise selection under VIF control.

### Feature selection

To combat the problem of p (the number of drug features plus cancer cell line features and interaction terms) ≫ n (the number of drug responses) in the training sets, we implemented Lasso [75] with Bayesian information criterion (BIC) [76] by an R package “HDeconometrics” [77] (the third component of our prediction framework in Fig 1) because Lasso is appropriate for estimating coefficients in high-dimensional space [78] and BIC provides an efficient approach to obtain the optimal lasso fit [79]. After feature selection, the remaining number of selected features for each cancer type is shown in Table 2.

### Neural network architecture

We built neural network models by using JMP^®^ [80]. We designed three types of neural network architectures in this study: single-layer, double-layer, and complex-double-layer. The numbers of hidden layer nodes follow the geometric pyramid rule [81]. Given N input nodes, there are ⌈*N* ^1*/*2^⌉ hidden nodes in a single-layer architecture. In a double-layer architecture, there are ⌈*N* ^2*/*3^⌉ and ⌈*N* ^1*/*3^⌉ hidden nodes in the first and second hidden layers, respectively. In a complex-double-layer architecture, there are *N* and ⌈*N* ^1*/*2^⌉ hidden nodes in the first and second hidden layers, respectively. The nodes among the two layers are fully connected. Biases are introduced into the input and hidden layers. The activation function of every node in the neural network is a hyperbolic tangent function (TanH). Newton’s method [82] is chosen as an optimizer by JMP.

To avoid overfitting, we implement 10-fold cross-validation, early stopping, and Lasso-style penalty function (absolute value penalty, i.e. L1 regularization [83]). Based on Occam’s razor principle [39], we started from a single-layer model for each cancer type. If the performance (average R^2^ of the intact validation sets across the 10 folds) is less than a threshold 0.8 in 200 iterations, we increased the iteration to 300; if the performance is still less than the threshold, we implemented a double-layer model for 200 iterations and so on until using a complex-double-layer model for 300 iterations. To increase the reproducibility of this study, fixed random seeds were assigned and all the codes for training and prediction models are available at https://github.com/leon1003/QSMART/.

### Other machine learning models

We compared neural networks with three other prediction algorithms with 10-fold cross-validation: random forests [84], support vector machine (SVM) [85], and Lasso [75]. Random forests were implemented by WEKA (version 3.8.3) [86] with default settings (“maxDepth” = 0, “bagSizePercent” = 100). For each cancer type, the number of iterations was decided based on the iterations used for each of the pre-trained neural network models (200 or 300 iterations) shown in S1 Table. SVM was implemented by the SMOreg function (SVM for regression) of WEKA with default kernel (“PolyKernel”) and optimizer (“RegSMOImproved”) settings. Lasso was implemented by an R package “glmnet” [87] with the default parameter setting for Lasso regression (alpha = 1 and family = “gaussian”). Additionally, we also compared our prediction models with two-way ANOVA analysis and MCA [35]. Because the purpose of two-way ANOVA analysis implemented by R was to quantify how much two factors (drug and cancer cell line) can explain the variation of drug response (adjusted R^2^ was used), the model used the drug and cancer cell line identifiers as inputs and did not undergo 10-fold cross-validation. The performance of MCA shown in Table 2 is based on its prediction for PKI response (details are available in S1 Data).

## Supporting information

**S1 Fig. Residual analyses for 23 cancer-centric models and the overall result of using QSMART with neural networks.**

**S2 Fig. Genome-wide mutational status (genomic fingerprints) across the 23 cancer types.**

**S3 Fig. Comparison between GDSC1 and GDSC2 in the GDSC release 8.0.**

**S4 Fig. Prediction performances of using QSMART model with neural networks for different PKI target groups.**

**S1 Table. The number of features at different feature levels and the prediction performance of neural networks.**

**S2 Table. Prediction performances of using genomic fingerprints.**

**S3 Table. Prediction performances of using no drug-mutation interaction terms.**

**S4 Table. Prediction performances of using no interaction terms.**

**S5 Table. Prediction performances of using random feature selection.**

**S6 Table. Prediction performances of using random 10X feature selection. S7 Table. Pathway enrichment analysis.**

**S8 Table. Drug-mutation relationships and their impact on IC**_**50**_ **in NSCLC cells.**

**S9 Table. Cancer cell line features.**

**S1 Data. MCA’s performance for PKI response prediction. S2 Data. Features for Lung (NSCLC) dataset.**

**S3 Data. Understudied proteins.**

**S4 Data. PKI target groups and PKI structures.**

## Acknowledgments

Funding for N.K. (R01GM114409 and U01CA239106) from the National Institutes of Health is acknowledged. Funding for P.M. (R01GM122080 and DMS-1903226) from NIH and NSF is acknowledged.

## Author Contributions

L.H., P.M., K.R., and N.K. designed the research. L.H. performed data integration. L.H., Y.W., H.C., and P.M. performed statistical analyses. L.H., Y.W., H.C., S.L., and K.R. performed machine learning methods. L.H., W.Y., Y.W., H.C., P.M., and N.K. analyzed the data and interpreted the results. L.H., W.Y., A.V., and N.K. wrote the manuscript. Y.W., H.C., S.L., P.M., and K.R. revised the manuscript. L.H. created the tables and figures. All authors reviewed the manuscript and approved the final version of the manuscript.

