## Supporting information for "Quantitative Structure-Mutation-Activity Relationship Tests (QSMART) Model for Protein Kinase Inhibitor Response Prediction"

#### Prediction performance for different PKI target groups

Some previous studies [1, 2] built both drug-centric and cancer-centric models to predict drug response. However, since the focus of our study is on investigating the drug-mutation relationships, we did not build drug-centric models. If we apply the framework in our study to a single drug, all the drug features will be the same across different drug response samples, and thus no significant drug features nor significant interaction terms will be captured. Moreover, if we built a drug-centric model for each combination of drug and cancer type, its sample size would not be enough to train a neural network model. Nevertheless, we were still interested in the prediction performances for different drugs. For each cancer type, the prediction model with the best validation performance across the 10-fold cross-validation was chosen as the final model. We used the model to predict drug response for the entire training set and then gathered prediction results for each PKI. Because the number of drug responses for a single drug in each cancer type dataset was small and not sufficient for seeing the overall performance, we pooled the results of PKIs according to their target groups shown in S4 Data into nine sets: AGC, CAMK, CK1, CMGC, STE, TK, TKL, Other, and Atypical. One PKI response prediction might be pooled in one or multiple sets since one PKI may have one or more drug targets classified as different protein kinase groups. We first analyzed the average actual $IC_{50}$ in different PKI target group sets. The result showed that if a drug inhibits a protein kinase classified as CMGC, CAMK, or AGC group, it has higher average $IC_{50}$ values for most of the cancer types (average $IC_{50}$ = 2.527, 2.389, and 2.331, respectively. S4 Fig panel a); if a drug inhibits Atypical and CK1 protein kinases, it has lower average $IC_{50}$ values (1.624 and 1.679, respectively). Note that this result was according to the data we collected from GDSC, and this might not be applied to all the cases. S4 Fig panel b and c show the detailed performances evaluated by $R^2$ and RMSE, respectively. To our surprise, we found the best is Atypical group ($R^2$ = 0.699 to 0.900 and overall $R^2$ = 0.871) and the worst is CAMK group ($R^2$ = 0.644 to 0.885 and overall $R^2$ = 0.786).

Even though atypical protein kinases lack canonical protein kinase domains, meaning their drug-target binding mode might not resemble canonical protein kinase's binding mode, the models could still predict atypical protein kinase inhibitor responses well. We speculated that the performance was supported by independent drug features, cancer cell line features, or the drug-mutation relationships from unknown off-targets. The mammalian target of rapamycin (mTOR), classified as Atypical group, is another potential factor to explain this result. mTOR regulates cell growth, proliferation, motility, and survival [3], and it is highly mutated in the cancer cell lines in our dataset: 67 out of 837 cell lines (8%) have mTOR mutations. Since it is critical to cell activity, the 6 drugs that inhibit mTOR (listed in S4 Data) might require less concentration to inhibit cancer cell line's activity and thus the average actual $IC_{50}$ is relatively low. Moreover, since mTOR is implicated in a broad category of pathways, each of its mutations provides more information about the sample's cancer cell line features to the prediction models and thus the overall prediction performance is relatively high. On the contrary, although the proteins in the CAMK group have canonical protein kinase domains, the models were not able to predict CAMK inhibitor responses well. We conjectured that this was because none of the PKIs in our dataset specifically inhibit CAMK group proteins, and thus the models were not tailored to capture CAMK inhibitor-specific features and interaction terms. Although there are 33 CAMK inhibitors in our dataset (S1 Data), all of them had at least one more target classified as other groups. Comparing to CAMK inhibitors, atypical PKIs had relatively higher specificity in this point of view: there are 29 atypical PKIs in our dataset and 8 of them (27.6%) only inhibit their targets classified as Atypical group.

### More explanations about the features in the NSCLC case study

Gene-level feature: "CNV_ROCK2_gain". This feature represents if Rho-associated protein kinase 2 (ROCK2) is either neutral or deleted in a cancer cell line (0, copy number losses) or amplified (1, copy number gains). ROCK2 is known to be essential for NSCLC's growth and invasion [4]. In the NSCLC dataset, ROCK2 is amplified in two cell lines: LC-1/sq and NCI-H1623; the source of the latter was from a patient with metastatic NSCLC. On average, the PKI responses involved in the cell lines with neutral or deleted ROCK2 showed lower $IC_{50}$ value than those with amplified ROCK2 (average actual $IC_{50} = 2.71$ vs. 3.49). By using the pre-trained model, however, we found that when the value of CNV_ROCK2_gain was replaced from 0 to 1 when other features were held constant, the estimated $IC_{50}$ decreased 0.14 on average (average predicted $IC_{50} = 2.71$ vs. 2.57). Although the coefficient of CNV_ROCK2_gain obtained from Lasso feature selection was 0.07, meaning it positively correlated to $IC_{50}$, the neural network model had not perfectly learned this trend.

Pathway-level feature: "REC_R_HSA_176298". This feature shows the number of mutations occurred in the proteins implicating in the reaction "Activation of claspin" (Reactome ID: R-HSA-176298). Claspin is an essential regulator for checkpoint kinase 1 (Chk1) activation, and it was found to be associated with regulating breast cancer proliferation [5, 6] and contributing to lung cancer radioresistance [7]. Interestingly, this feature was also selected in our PKI response prediction model for breast cancer cell lines. On average, the NSCLC cell lines without mutations related to activation of claspin had lower PKI responses than those with mutations related to this reaction (average actual $IC_{50} = 2.66$ vs. 3.27). Based on the pre-trained neural network model and our NSCLC dataset, every unit increase in REC_R_HSA_176298 is associated with a 0.52 unit increase in $IC_{50}$ on average (average predicted $IC_{50} = 2.73$ vs. 3.25).


**S1 Table . The number of features at different feature levels and the prediction performance of neural networks.**

| Cancer type | #$IC_{50}$ | #All Features | #Drug Features | #Cancer cell line features | | | | | | | #Interaction terms | | | | | #Nodes | | #Tours | Performance | | |
|---|---|---|---|---|---|---|---|---|---|---|---|---|---|---|---|---|---|---|---|---|---|
| | | | | Residue | Motif | Domain | Gene | Family | Pathway | Sample | DxM | PPI | RECx | PWYx | GOx | 1st | 2nd | | RMSE | $R^2$ | AUC |
| AG | 2,971 | 62 | 38 | 0 | 0 | 0 | 4 | 1 | 0 | 0 | 9 | 9 | 0 | 1 | 0 | 38 | 8 | 200 | 0.851 | 0.815 | 0.976 |
| Bone | 3,410 | 84 | 52 | 0 | 1 | 0 | 1 | 0 | 11 | 0 | 4 | 11 | 0 | 3 | 1 | 10 | 0 | 300 | 0.812 | 0.856 | 0.984 |
| Breast | 4,706 | 129 | 70 | 5 | 0 | 1 | 10 | 0 | 15 | 0 | 12 | 6 | 1 | 5 | 4 | 26 | 6 | 200 | 0.714 | 0.880 | 0.986 |
| CNS | 4,250 | 114 | 65 | 0 | 0 | 0 | 9 | 1 | 12 | 1 | 11 | 6 | 1 | 4 | 4 | 11 | 0 | 300 | 0.785 | 0.858 | 0.980 |
| Cervix | 1,044 | 37 | 29 | 0 | 0 | 0 | 2 | 0 | 1 | 0 | 1 | 4 | 0 | 0 | 0 | 7 | 0 | 200 | 0.770 | 0.864 | 0.989 |
| Endometrium | 1,073 | 33 | 21 | 0 | 0 | 0 | 0 | 0 | 3 | 1 | 4 | 3 | 0 | 0 | 1 | 11 | 4 | 200 | 0.733 | 0.878 | 0.982 |
| Haematopoietic | 4,204 | 119 | 58 | 3 | 0 | 2 | 9 | 0 | 13 | 0 | 28 | 2 | 0 | 0 | 4 | 11 | 0 | 200 | 0.906 | 0.858 | 0.971 |
| Kidney | 2,458 | 73 | 51 | 0 | 0 | 0 | 1 | 0 | 1 | 1 | 0 | 17 | 1 | 0 | 1 | 9 | 0 | 200 | 0.877 | 0.836 | 0.986 |
| Large intestine | 4,628 | 141 | 53 | 10 | 1 | 1 | 4 | 0 | 8 | 0 | 50 | 10 | 1 | 3 | 0 | 12 | 0 | 300 | 0.923 | 0.814 | 0.974 |
| Liver | 1,348 | 48 | 35 | 0 | 0 | 0 | 0 | 2 | 2 | 0 | 2 | 6 | 0 | 0 | 1 | 7 | 0 | 200 | 0.844 | 0.836 | 0.985 |
| Lung (NSCLC) | 9,205 | 207 | 72 | 7 | 0 | 0 | 9 | 4 | 21 | 1 | 47 | 27 | 1 | 3 | 15 | 15 | 0 | 200 | 0.809 | 0.854 | 0.982 |
| Lung (others) | 7,206 | 162 | 58 | 2 | 0 | 0 | 3 | 1 | 11 | 1 | 46 | 23 | 0 | 4 | 13 | 30 | 6 | 200 | 0.756 | 0.859 | 0.983 |
| Lymphoid | 13,302 | 291 | 72 | 54 | 0 | 2 | 11 | 1 | 14 | 2 | 86 | 39 | 4 | 0 | 6 | 18 | 0 | 300 | 0.757 | 0.873 | 0.980 |
| Oesophagus | 3,337 | 91 | 58 | 0 | 0 | 0 | 8 | 0 | 9 | 0 | 4 | 9 | 0 | 1 | 2 | 10 | 0 | 200 | 0.857 | 0.841 | 0.972 |
| Ovary | 3,502 | 113 | 64 | 2 | 0 | 1 | 9 | 3 | 5 | 0 | 9 | 17 | 1 | 0 | 2 | 11 | 0 | 200 | 0.867 | 0.844 | 0.987 |
| Pancreas | 2,421 | 84 | 60 | 0 | 0 | 1 | 2 | 1 | 3 | 0 | 0 | 13 | 0 | 3 | 1 | 10 | 0 | 200 | 0.877 | 0.833 | 0.990 |
| Pleura | 1,431 | 36 | 23 | 0 | 1 | 2 | 2 | 0 | 0 | 0 | 0 | 8 | 0 | 0 | 0 | 11 | 4 | 200 | 0.894 | 0.805 | 0.966 |
| Skin | 5,732 | 132 | 64 | 9 | 0 | 1 | 7 | 0 | 13 | 0 | 15 | 15 | 0 | 3 | 5 | 12 | 0 | 200 | 0.810 | 0.875 | 0.987 |
| Soft tissue | 1,938 | 63 | 45 | 0 | 1 | 0 | 1 | 1 | 7 | 0 | 2 | 5 | 0 | 1 | 0 | 8 | 0 | 200 | 0.941 | 0.818 | 0.975 |
| Stomach | 2,327 | 83 | 49 | 0 | 0 | 0 | 8 | 1 | 4 | 0 | 16 | 5 | 0 | 0 | 0 | 20 | 5 | 200 | 0.837 | 0.836 | 0.981 |
| Thyroid | 1,352 | 33 | 25 | 0 | 0 | 0 | 5 | 0 | 0 | 0 | 0 | 2 | 0 | 1 | 0 | 6 | 0 | 300 | 0.963 | 0.830 | 0.973 |
| UAT | 3,856 | 126 | 74 | 1 | 0 | 0 | 4 | 1 | 8 | 0 | 13 | 18 | 0 | 2 | 5 | 12 | 0 | 300 | 0.798 | 0.869 | 0.986 |
| Urinary tract | 1,454 | 68 | 47 | 0 | 0 | 0 | 3 | 0 | 2 | 0 | 9 | 6 | 0 | 0 | 1 | 9 | 0 | 200 | 0.750 | 0.863 | 0.988 |
| Overall | 87,155 | | | | | | | | | | | | | | | | | | 0.818 | 0.861 | 0.981 |

AG: autonomic ganglia; AUC: area under the ROC Curve; CNS: central nervous system; DxM: drug-mutation interaction term; GOx: biological process interaction; NSCLC: non-small cell lung cancer; PPI: protein-protein interaction; PWYx: pathway-pathway interaction; $R^2$: coefficient of determination; RECx: reaction-reaction interaction; RMSE: root-mean-square error; UAT: upper aerodigestive tract; #$IC_{50}$: the number of drug responses; #Nodes: the number of nodes in the first and second hidden layers of neural networks; #Tours: the number of times to restart the fitting process.

**S2 Table . Prediction performances of using genomic fingerprints.**

| Cancer type | $\#IC_{50}$ | #All Features | #Drug Features | #Genomics Fingerprints | #Nodes | | Performance ($R^2$) | | | |
|---|---|---|---|---|---|---|---|---|---|---|
| | | | | | 1st | 2nd | NN | RF | SVM | Lasso |
| AG | 2,971 | 62 | 60 | 2 | 38 | 8 | **0.696** | 0.603 | 0.555 | 0.580 |
| Bone | 3,410 | 84 | 50 | 34 | 10 | 0 | **0.630** | 0.614 | 0.543 | 0.572 |
| Breast | 4,706 | 129 | 68 | 61 | 26 | 6 | **0.732** | 0.628 | 0.602 | 0.620 |
| CNS | 4,250 | 114 | 77 | 37 | 11 | 0 | **0.769** | 0.701 | 0.678 | 0.698 |
| Cervix | 1,044 | 37 | 16 | 21 | 7 | 0 | **0.663** | 0.586 | 0.556 | 0.574 |
| Endometrium | 1,073 | 33 | 32 | 1 | 11 | 4 | **0.735** | 0.634 | 0.517 | 0.553 |
| Haematopoietic | 4,204 | 119 | 64 | 55 | 11 | 0 | **0.694** | 0.622 | 0.567 | 0.596 |
| Kidney | 2,458 | 73 | 54 | 19 | 9 | 0 | **0.771** | 0.701 | 0.596 | 0.627 |
| Large intestine | 4,628 | 141 | 73 | 68 | 12 | 0 | **0.703** | 0.671 | 0.658 | 0.677 |
| Liver | 1,348 | 48 | 33 | 15 | 7 | 0 | **0.787** | 0.631 | 0.537 | 0.577 |
| Lung (NSCLC) | 9,205 | 207 | 73 | 134 | 15 | 0 | **0.671** | 0.640 | 0.612 | 0.626 |
| Lung (others) | 7,206 | 162 | 66 | 96 | 30 | 6 | **0.735** | 0.641 | 0.601 | 0.616 |
| Lymphoid | 13,302 | 291 | 92 | 199 | 18 | 0 | **0.812** | 0.687 | 0.705 | 0.717 |
| Oesophagus | 3,337 | 91 | 66 | 25 | 10 | 0 | 0.693 | **0.699** | 0.638 | 0.662 |
| Ovary | 3,502 | 113 | 60 | 53 | 11 | 0 | **0.759** | 0.673 | 0.636 | 0.657 |
| Pancreas | 2,421 | 84 | 57 | 27 | 10 | 0 | **0.770** | 0.687 | 0.669 | 0.697 |
| Pleura | 1,431 | 36 | 33 | 3 | 11 | 4 | **0.787** | 0.663 | 0.515 | 0.550 |
| Skin | 5,732 | 132 | 82 | 50 | 12 | 0 | **0.730** | 0.704 | 0.673 | 0.693 |
| Soft tissue | 1,938 | 63 | 53 | 10 | 8 | 0 | **0.773** | 0.677 | 0.604 | 0.644 |
| Stomach | 2,327 | 83 | 63 | 20 | 20 | 5 | **0.728** | 0.627 | 0.598 | 0.625 |
| Thyroid | 1,352 | 33 | 27 | 6 | 6 | 0 | **0.788** | 0.648 | 0.526 | 0.564 |
| UAT | 3,856 | 126 | 73 | 53 | 12 | 0 | **0.807** | 0.737 | 0.726 | 0.744 |
| Urinary tract | 1,454 | 68 | 55 | 13 | 9 | 0 | **0.799** | 0.641 | 0.648 | 0.681 |
| Overall | 87,155 | | | | | | **0.752** | 0.680 | 0.646 | 0.655 |

The best performance for each cancer type is highlighted in bold. AG: autonomic ganglia; CNS: central nervous system; NN: neural networks; NSCLC: non-small cell lung cancer; $R^2$: coefficient of determination; RF: random forests; SVM: support vector machine; UAT: upper aerodigestive tract; $\#IC_{50}$: the number of drug responses; #Nodes: the number of nodes in the first and second hidden layers of neural networks.

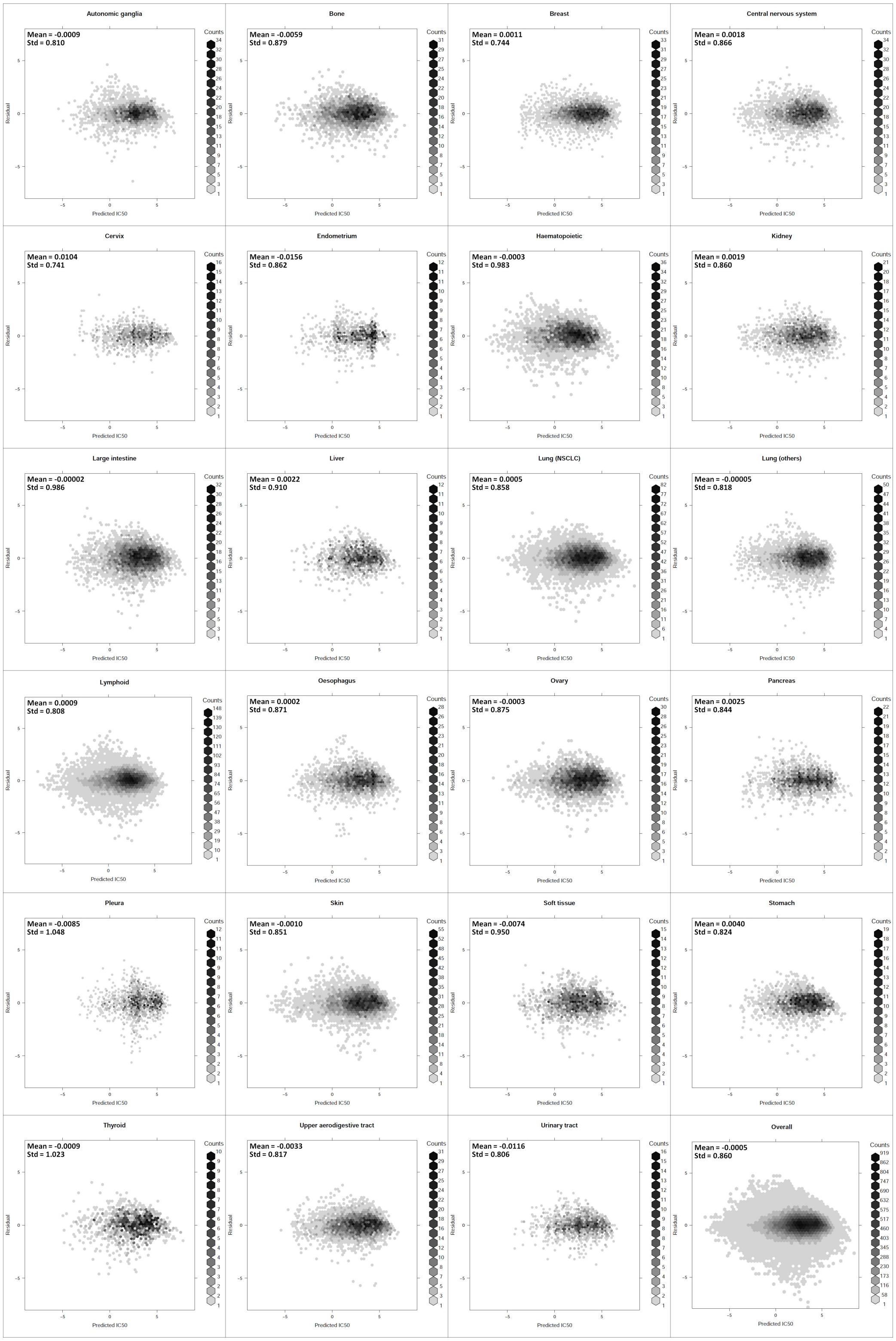


**S1 Fig . Residual analyses for 23 cancer-centric models and the overall result of using QSMART with neural networks.** X-axis: predicted $IC_{50}$; y-axis: residuals, defined as actual $IC_{50}$ minus predicted $IC_{50}$. Residuals mean and standard deviation are shown for each cancer type.

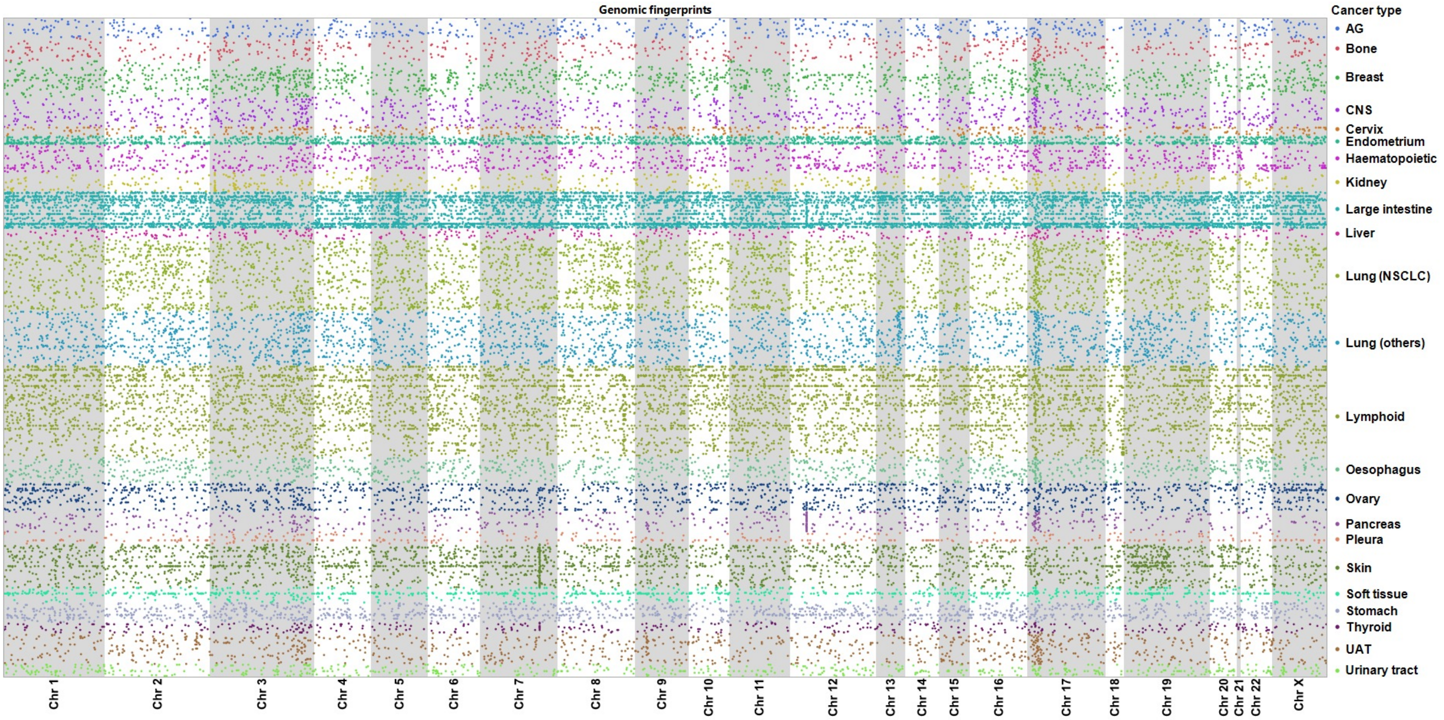


**S2 Fig . Genome-wide mutational status (genomic fingerprints) across all 23 cancer types.** AG: autonomic ganglia; CNS: central nervous system; NSCLC: non-small cell lung cancer; UAT: upper aerodigestive tract.

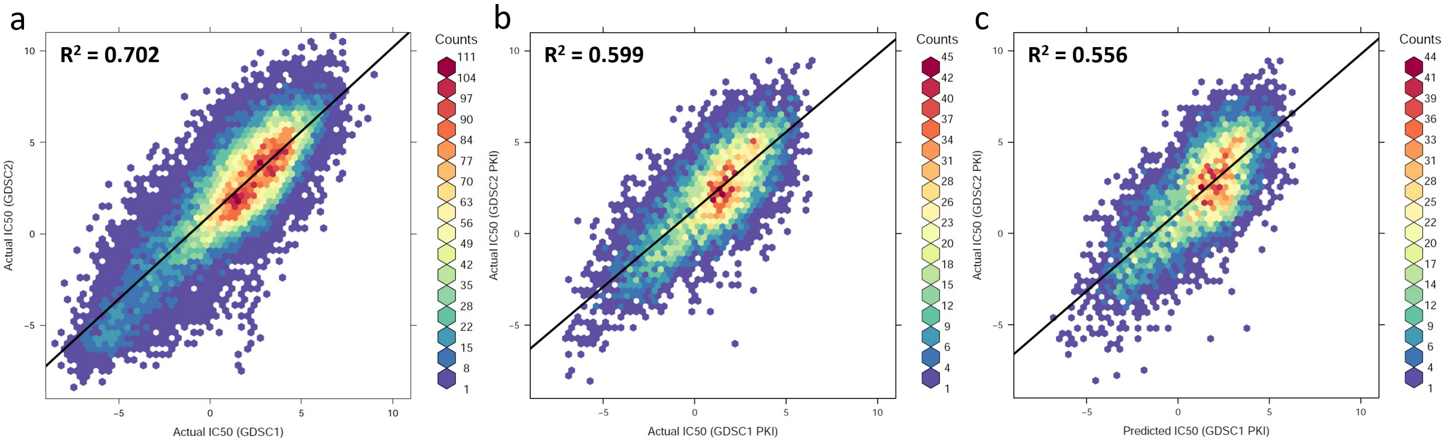


**S3 Fig . Comparison between GDSC1 and GDSC2 in the GDSC release 8.0.** GDSC1 (the old drug response dataset) and GDSC2 (the new drug response dataset) were generated based on different types of assays. Cell viability was measured using either Resazurin or Syto60 in GDSC1, while it was measured based on Promega CellTiter-Glo® in GDSC2. In total, there are 22,624 drug-cancer cell line pairs found in both datasets; the experiments of all these pairs were done by Wellcome Sanger Institute. (a) The hexbin plot shows the actual $IC_{50}$ from GDSC1 (x-axis) versus the actual $IC_{50}$ from GDSC2 (y-axis); a fitted regression line and its $R^2$ are shown. (b) There are 7,283 PKI-cancer cell line pairs found in both GDSC1 and GDSC2. The hexbin plot shows the PKI's actual $IC_{50}$ from GDSC1 (x-axis) versus the PKI's actual $IC_{50}$ from GDSC2 (y-axis). (c) Based on the prediction result of our QSMART with neural network models trained by GDSC1 data, the hexbin plot shows the PKI's predicted $IC_{50}$ (x-axis) versus the PKI's actual $IC_{50}$ from GDSC2 (y-axis).

**a** Average $IC_{50}$

| Cancer type | PKI target group | | | | | | | | |
|---|---|---|---|---|---|---|---|---|---|
| | AGC | CAMK | CK1 | CMGC | STE | TK | TKL | Other | Atypical |
| Autonomic ganglia | 2.104 | 2.256 | 1.848 | 2.495 | 2.006 | 2.123 | 1.984 | 1.941 | 1.469 |
| Bone | 2.036 | 2.120 | 1.389 | 2.249 | 1.887 | 1.914 | 1.786 | 1.727 | 1.296 |
| Breast | 2.953 | 3.027 | 2.156 | 3.081 | 2.714 | 2.721 | 2.692 | 2.761 | 2.316 |
| Central nervous system | 2.434 | 2.486 | 1.901 | 2.658 | 2.187 | 2.339 | 2.178 | 2.282 | 1.802 |
| Cervix | 3.120 | 3.068 | 2.384 | 3.231 | 2.709 | 2.644 | 2.964 | 2.852 | 2.181 |
| Endometrium | 2.479 | 2.585 | 1.496 | 2.681 | 2.285 | 2.374 | 2.259 | 2.296 | 1.855 |
| Haematopoietic | 1.098 | 1.181 | 0.694 | 1.344 | 0.880 | 1.049 | 0.976 | 0.816 | 0.226 |
| Kidney | 2.410 | 2.612 | 2.061 | 2.770 | 2.238 | 2.327 | 2.284 | 2.403 | 1.825 |
| Large intestine | 2.820 | 2.868 | 2.133 | 3.008 | 2.355 | 2.601 | 2.624 | 2.571 | 2.192 |
| Liver | 2.844 | 2.910 | 2.495 | 2.999 | 2.461 | 2.452 | 2.707 | 2.634 | 2.045 |
| Lung (NSCLC) | 2.806 | 2.827 | 2.070 | 2.893 | 2.461 | 2.526 | 2.609 | 2.541 | 2.128 |
| Lung (others) | 2.589 | 2.676 | 1.905 | 2.818 | 2.485 | 2.488 | 2.499 | 2.299 | 1.951 |
| Lymphoid | 1.301 | 1.441 | 0.726 | 1.565 | 1.285 | 1.458 | 1.263 | 1.069 | 0.492 |
| Oesophagus | 2.480 | 2.589 | 1.755 | 2.702 | 2.216 | 2.285 | 2.379 | 2.335 | 1.919 |
| Ovary | 2.806 | 2.679 | 1.909 | 2.851 | 2.225 | 2.473 | 2.462 | 2.446 | 1.935 |
| Pancreas | 3.051 | 3.034 | 2.272 | 3.050 | 2.483 | 2.657 | 2.801 | 2.667 | 2.257 |
| Pleura | 2.976 | 3.050 | 2.619 | 3.168 | 2.768 | 2.782 | 2.820 | 2.911 | 2.338 |
| Skin | 2.419 | 2.426 | 1.724 | 2.686 | 1.853 | 2.311 | 2.025 | 2.285 | 1.680 |
| Soft tissue | 2.204 | 2.293 | 1.607 | 2.455 | 2.001 | 2.126 | 1.990 | 2.043 | 1.365 |
| Stomach | 2.605 | 2.707 | 1.857 | 2.750 | 2.297 | 2.408 | 2.446 | 2.437 | 1.821 |
| Thyroid | 2.482 | 2.497 | 2.009 | 2.662 | 2.024 | 2.267 | 2.141 | 2.397 | 1.716 |
| Upper aerodigestive tract | 2.744 | 2.608 | 2.005 | 2.771 | 2.198 | 2.330 | 2.511 | 2.435 | 1.914 |
| Urinary tract | 2.833 | 2.776 | 2.307 | 2.869 | 2.410 | 2.469 | 2.601 | 2.544 | 2.048 |
| Overall | 2.331 | 2.389 | 1.679 | 2.527 | 2.055 | 2.199 | 2.144 | 2.105 | 1.624 |

Max / Min

**b** Overall $R^2$

| Cancer type | PKI target group | | | | | | | | |
|---|---|---|---|---|---|---|---|---|---|
| | AGC | CAMK | CK1 | CMGC | STE | TK | TKL | Other | Atypical |
| Autonomic ganglia | 0.861 | 0.782 | 0.865 | 0.845 | 0.842 | 0.806 | 0.822 | 0.826 | 0.866 |
| Bone | 0.835 | 0.775 | 0.850 | 0.846 | 0.797 | 0.810 | 0.811 | 0.826 | 0.856 |
| Breast | 0.874 | 0.805 | 0.888 | 0.870 | 0.861 | 0.854 | 0.845 | 0.867 | 0.893 |
| Central nervous system | 0.817 | 0.710 | 0.856 | 0.823 | 0.777 | 0.802 | 0.785 | 0.799 | 0.860 |
| Cervix | 0.891 | 0.885 | 0.927 | 0.892 | 0.891 | 0.868 | 0.859 | 0.898 | 0.900 |
| Endometrium | 0.842 | 0.806 | 0.892 | 0.867 | 0.829 | 0.815 | 0.857 | 0.826 | 0.895 |
| Haematopoietic | 0.837 | 0.804 | 0.872 | 0.846 | 0.803 | 0.828 | 0.827 | 0.827 | 0.847 |
| Kidney | 0.848 | 0.770 | 0.901 | 0.856 | 0.824 | 0.822 | 0.853 | 0.828 | 0.891 |
| Large intestine | 0.773 | 0.644 | 0.732 | 0.763 | 0.771 | 0.746 | 0.739 | 0.757 | 0.785 |
| Liver | 0.803 | 0.708 | 0.831 | 0.785 | 0.841 | 0.808 | 0.780 | 0.829 | 0.820 |
| Lung (NSCLC) | 0.821 | 0.741 | 0.840 | 0.825 | 0.821 | 0.817 | 0.799 | 0.820 | 0.857 |
| Lung (others) | 0.841 | 0.776 | 0.876 | 0.845 | 0.824 | 0.817 | 0.817 | 0.833 | 0.871 |
| Lymphoid | 0.857 | 0.812 | 0.874 | 0.869 | 0.823 | 0.837 | 0.830 | 0.841 | 0.873 |
| Oesophagus | 0.829 | 0.689 | 0.841 | 0.846 | 0.785 | 0.807 | 0.825 | 0.800 | 0.873 |
| Ovary | 0.825 | 0.784 | 0.865 | 0.822 | 0.857 | 0.815 | 0.811 | 0.848 | 0.856 |
| Pancreas | 0.837 | 0.750 | 0.825 | 0.814 | 0.855 | 0.840 | 0.806 | 0.863 | 0.843 |
| Pleura | 0.670 | 0.709 | 0.794 | 0.691 | 0.699 | 0.680 | 0.769 | 0.642 | 0.699 |
| Skin | 0.841 | 0.772 | 0.834 | 0.844 | 0.868 | 0.826 | 0.837 | 0.823 | 0.871 |
| Soft tissue | 0.840 | 0.732 | 0.877 | 0.821 | 0.786 | 0.795 | 0.810 | 0.800 | 0.878 |
| Stomach | 0.832 | 0.756 | 0.850 | 0.835 | 0.815 | 0.818 | 0.811 | 0.811 | 0.886 |
| Thyroid | 0.822 | 0.756 | 0.806 | 0.801 | 0.824 | 0.762 | 0.776 | 0.801 | 0.809 |
| Upper aerodigestive tract | 0.827 | 0.791 | 0.870 | 0.841 | 0.859 | 0.852 | 0.817 | 0.847 | 0.881 |
| Urinary tract | 0.839 | 0.764 | 0.850 | 0.832 | 0.859 | 0.824 | 0.813 | 0.833 | 0.859 |
| Overall | 0.846 | 0.786 | 0.863 | 0.849 | 0.832 | 0.826 | 0.828 | 0.837 | 0.871 |

Max / Min

**c** Overall RMSE

| Cancer type | PKI target group | | | | | | | | |
|---|---|---|---|---|---|---|---|---|---|
| | AGC | CAMK | CK1 | CMGC | STE | TK | TKL | Other | Atypical |
| Autonomic ganglia | 0.825 | 0.877 | 0.794 | 0.749 | 0.841 | 0.855 | 0.786 | 0.879 | 0.806 |
| Bone | 0.913 | 0.914 | 0.898 | 0.808 | 0.944 | 0.907 | 0.854 | 0.940 | 0.865 |
| Breast | 0.766 | 0.800 | 0.776 | 0.713 | 0.787 | 0.760 | 0.770 | 0.802 | 0.748 |
| Central nervous system | 0.966 | 1.034 | 0.920 | 0.858 | 1.017 | 0.888 | 0.932 | 0.975 | 0.890 |
| Cervix | 0.693 | 0.663 | 0.696 | 0.656 | 0.765 | 0.733 | 0.772 | 0.700 | 0.748 |
| Endometrium | 0.901 | 0.866 | 0.785 | 0.756 | 0.884 | 0.885 | 0.799 | 0.932 | 0.788 |
| Haematopoietic | 0.962 | 0.991 | 0.909 | 0.916 | 1.054 | 0.993 | 0.954 | 1.000 | 0.893 |
| Kidney | 0.895 | 0.971 | 0.736 | 0.786 | 0.935 | 0.875 | 0.801 | 0.917 | 0.800 |
| Large intestine | 1.018 | 1.077 | 1.070 | 0.938 | 1.112 | 1.007 | 0.976 | 1.029 | 0.988 |
| Liver | 0.877 | 0.935 | 0.809 | 0.876 | 0.848 | 0.876 | 0.864 | 0.832 | 0.960 |
| Lung (NSCLC) | 0.901 | 0.932 | 0.883 | 0.823 | 0.923 | 0.868 | 0.868 | 0.901 | 0.840 |
| Lung (others) | 0.865 | 0.889 | 0.778 | 0.765 | 0.877 | 0.831 | 0.783 | 0.892 | 0.791 |
| Lymphoid | 0.860 | 0.889 | 0.853 | 0.783 | 0.873 | 0.821 | 0.834 | 0.902 | 0.798 |
| Oesophagus | 0.937 | 1.066 | 0.865 | 0.776 | 1.017 | 0.921 | 0.809 | 0.986 | 0.832 |
| Ovary | 0.884 | 0.908 | 0.801 | 0.835 | 0.866 | 0.889 | 0.867 | 0.847 | 0.871 |
| Pancreas | 0.843 | 0.911 | 0.969 | 0.847 | 0.910 | 0.810 | 0.860 | 0.820 | 0.875 |
| Pleura | 1.181 | 0.924 | 0.944 | 1.054 | 1.133 | 1.098 | 0.925 | 1.179 | 1.276 |
| Skin | 0.958 | 1.028 | 0.935 | 0.830 | 0.992 | 0.882 | 0.927 | 0.972 | 0.905 |
| Soft tissue | 0.987 | 1.072 | 0.886 | 0.914 | 1.064 | 0.979 | 0.954 | 1.041 | 0.862 |
| Stomach | 0.873 | 0.879 | 0.831 | 0.777 | 0.912 | 0.834 | 0.815 | 0.903 | 0.772 |
| Thyroid | 1.004 | 1.047 | 1.003 | 0.947 | 1.088 | 1.074 | 1.086 | 1.022 | 1.120 |
| Upper aerodigestive tract | 0.855 | 0.836 | 0.786 | 0.768 | 0.864 | 0.810 | 0.816 | 0.834 | 0.783 |
| Urinary tract | 0.794 | 0.808 | 0.797 | 0.743 | 0.770 | 0.810 | 0.806 | 0.842 | 0.808 |
| Overall | 0.901 | 0.936 | 0.868 | 0.818 | 0.934 | 0.877 | 0.862 | 0.922 | 0.855 |

Max / Min

**S4 Fig . Prediction performances of using QSMART model with neural networks for different PKI target groups.** (a) Average actual $IC_{50}$ of different PKI target groups across 23 cancer types. (b) The prediction performances (in $R^2$) of using QSMART model with neural networks for different PKI target groups. (c) The prediction performances (in RMSE: root-mean-square error) of using QSMART model with neural networks for different PKI target groups. NSCLC: non-small cell lung cancer.

**S3 Table . Prediction performances of using no drug-mutation interaction terms.**

| Cancer type | $\#IC_{50}$ | #All Features | #Drug Features | #Cancer features | | #Interaction terms | | | | #Nodes | | Performance ($R^2$) | | | |
|---|---|---|---|---|---|---|---|---|---|---|---|---|---|---|---|
| | | | | Residue | Others | PPI | RECx | PWYx | GOx | 1st | 2nd | NN | RF | SVM | Lasso |
| AG | 2,971 | 62 | 44 | 0 | 6 | 10 | 0 | 1 | 1 | 38 | 8 | **0.790** | 0.373 | 0.300 | 0.338 |
| Bone | 3,410 | 84 | 58 | 0 | 13 | 11 | 0 | 1 | 1 | 10 | 0 | **0.836** | 0.472 | 0.359 | 0.404 |
| Breast | 4,706 | 129 | 74 | 3 | 31 | 6 | 1 | 9 | 5 | 26 | 6 | **0.880** | 0.546 | 0.444 | 0.490 |
| CNS | 4,250 | 114 | 74 | 0 | 28 | 2 | 0 | 5 | 5 | 11 | 0 | **0.867** | 0.604 | 0.394 | 0.464 |
| Cervix | 1,044 | 37 | 31 | 0 | 2 | 4 | 0 | 0 | 0 | 7 | 0 | **0.891** | 0.479 | 0.445 | 0.492 |
| Endometrium | 1,073 | 33 | 25 | 0 | 5 | 3 | 0 | 0 | 0 | 11 | 4 | **0.807** | 0.323 | 0.314 | 0.352 |
| Haematopoietic | 4,204 | 119 | 76 | 4 | 27 | 3 | 0 | 0 | 9 | 11 | 0 | **0.861** | 0.572 | 0.37 | 0.436 |
| Kidney | 2,458 | 73 | 53 | 0 | 4 | 15 | 0 | 0 | 1 | 9 | 0 | **0.750** | 0.489 | 0.405 | 0.460 |
| Large intestine | 4,628 | 141 | 76 | 20 | 25 | 11 | 7 | 2 | 0 | 12 | 0 | **0.837** | 0.520 | 0.453 | 0.500 |
| Liver | 1,348 | 48 | 35 | 0 | 5 | 7 | 0 | 0 | 1 | 7 | 0 | **0.777** | 0.486 | 0.398 | 0.450 |
| Lung (NSCLC) | 9,205 | 207 | 80 | 36 | 36 | 26 | 0 | 9 | 20 | 15 | 0 | **0.726** | 0.507 | 0.467 | 0.511 |
| Lung (others) | 7,206 | 162 | 80 | 12 | 21 | 27 | 0 | 9 | 13 | 30 | 6 | **0.892** | 0.465 | 0.419 | 0.469 |
| Lymphoid | 13,302 | 291 | 80 | 123 | 34 | 45 | 5 | 0 | 4 | 18 | 0 | **0.892** | 0.455 | 0.434 | 0.485 |
| Oesophagus | 3,337 | 91 | 64 | 0 | 14 | 10 | 0 | 2 | 1 | 10 | 0 | **0.830** | 0.523 | 0.415 | 0.479 |
| Ovary | 3,502 | 113 | 69 | 2 | 14 | 18 | 2 | 6 | 2 | 11 | 0 | **0.850** | 0.536 | 0.477 | 0.533 |
| Pancreas | 2,421 | 84 | 60 | 0 | 7 | 13 | 0 | 3 | 1 | 10 | 0 | **0.839** | 0.539 | 0.505 | 0.562 |
| Pleura | 1,431 | 36 | 25 | 0 | 4 | 7 | 0 | 0 | 0 | 11 | 4 | **0.701** | 0.275 | 0.321 | 0.363 |
| Skin | 5,732 | 132 | 63 | 15 | 28 | 10 | 2 | 7 | 7 | 12 | 0 | **0.864** | 0.597 | 0.413 | 0.454 |
| Soft tissue | 1,938 | 63 | 46 | 0 | 11 | 5 | 0 | 1 | 0 | 8 | 0 | **0.728** | 0.504 | 0.401 | 0.459 |
| Stomach | 2,327 | 83 | 58 | 2 | 18 | 4 | 0 | 0 | 1 | 20 | 5 | **0.874** | 0.523 | 0.395 | 0.446 |
| Thyroid | 1,352 | 33 | 25 | 0 | 5 | 2 | 0 | 1 | 0 | 6 | 0 | **0.653** | 0.403 | 0.396 | 0.435 |
| UAT | 3,856 | 126 | 80 | 3 | 17 | 18 | 0 | 1 | 7 | 12 | 0 | **0.779** | 0.651 | 0.534 | 0.597 |
| Urinary tract | 1,454 | 68 | 54 | 0 | 5 | 9 | 0 | 0 | 0 | 9 | 0 | **0.854** | 0.513 | 0.466 | 0.523 |
| Overall | 87,155 | | | | | | | | | | | **0.839** | 0.506 | 0.444 | 0.477 |

The best performance for each cancer type is highlighted in bold. AG: autonomic ganglia; CNS: central nervous system; PPI: protein-protein interaction; GOx: biological process interaction; NSCLC: non-small cell lung cancer; PWYx: pathway-pathway interaction; $R^2$: coefficient of determination; RECx: reaction-reaction interaction; UAT: upper aerodigestive tract; #IC50: the number of drug responses; #Nodes: the number of nodes in the first and second hidden layers of neural networks.

**S4 Table . Prediction performances of using no interaction terms.**

| Cancer type | $\#IC_{50}$ | #All Features | #Drug Features | #Cancer features | | #Nodes | | Performance ($R^2$) | | | |
|---|---|---|---|---|---|---|---|---|---|---|---|
| | | | | Residue | Others | 1st | 2nd | NN | RF | SVM | Lasso |
| AG | 2,971 | 62 | 46 | 0 | 16 | 38 | 8 | **0.833** | 0.425 | 0.298 | 0.341 |
| Bone | 3,410 | 84 | 59 | 0 | 25 | 10 | 0 | **0.692** | 0.501 | 0.356 | 0.406 |
| Breast | 4,706 | 129 | 77 | 9 | 43 | 26 | 6 | **0.901** | 0.543 | 0.443 | 0.492 |
| CNS | 4,250 | 114 | 72 | 0 | 42 | 11 | 0 | **0.745** | 0.576 | 0.394 | 0.463 |
| Cervix | 1,044 | 37 | 29 | 0 | 8 | 7 | 0 | **0.867** | 0.453 | 0.436 | 0.481 |
| Endometrium | 1,073 | 33 | 25 | 0 | 8 | 11 | 4 | **0.698** | 0.326 | 0.315 | 0.352 |
| Haematopoietic | 4,204 | 119 | 71 | 14 | 34 | 11 | 0 | **0.877** | 0.549 | 0.373 | 0.431 |
| Kidney | 2,458 | 73 | 47 | 5 | 21 | 9 | 0 | **0.783** | 0.486 | 0.396 | 0.447 |
| Large intestine | 4,628 | 141 | 73 | 38 | 30 | 12 | 0 | **0.732** | 0.565 | 0.453 | 0.499 |
| Liver | 1,348 | 48 | 38 | 1 | 9 | 7 | 0 | **0.842** | 0.549 | 0.397 | 0.461 |
| Lung (NSCLC) | 9,205 | 207 | 80 | 31 | 96 | 15 | 0 | **0.831** | 0.559 | 0.466 | 0.511 |
| Lung (others) | 7,206 | 162 | 78 | 6 | 78 | 30 | 6 | **0.898** | 0.536 | 0.420 | 0.469 |
| Lymphoid | 13,302 | 291 | 80 | 116 | 95 | 18 | 0 | **0.814** | 0.569 | 0.434 | 0.485 |
| Oesophagus | 3,337 | 91 | 59 | 0 | 32 | 10 | 0 | **0.745** | 0.486 | 0.416 | 0.472 |
| Ovary | 3,502 | 113 | 75 | 4 | 34 | 11 | 0 | **0.701** | 0.637 | 0.473 | 0.538 |
| Pancreas | 2,421 | 84 | 63 | 0 | 21 | 10 | 0 | **0.856** | 0.607 | 0.519 | 0.571 |
| Pleura | 1,431 | 36 | 25 | 0 | 11 | 11 | 4 | **0.824** | 0.290 | 0.320 | 0.363 |
| Skin | 5,732 | 132 | 68 | 11 | 53 | 12 | 0 | **0.887** | 0.614 | 0.408 | 0.457 |
| Soft tissue | 1,938 | 63 | 44 | 0 | 19 | 8 | 0 | **0.816** | 0.486 | 0.399 | 0.454 |
| Stomach | 2,327 | 83 | 52 | 5 | 26 | 20 | 5 | **0.783** | 0.445 | 0.386 | 0.434 |
| Thyroid | 1,352 | 33 | 27 | 0 | 6 | 6 | 0 | **0.696** | 0.434 | 0.436 | 0.465 |
| UAT | 3,856 | 126 | 75 | 6 | 45 | 12 | 0 | **0.912** | 0.692 | 0.538 | 0.595 |
| Urinary tract | 1,454 | 68 | 54 | 1 | 13 | 9 | 0 | **0.581** | 0.556 | 0.466 | 0.523 |
| Overall | 87,155 | | | | | | | **0.823** | 0.541 | 0.444 | 0.477 |

The best performance for each cancer type is highlighted in bold. AG: autonomic ganglia; CNS: central nervous system; NN: neural networks; NSCLC: non-small cell lung cancer; $R^2$: coefficient of determination; RF: random forests; SVM: support vector machine; UAT: upper aerodigestive tract; $\#IC_{50}$: the number of drug responses; #Nodes: the number of nodes in the first and second hidden layers of neural networks.

**S5 Table . Prediction performances of using random feature selection.**

| Cancer type | $\#IC_{50}$ | #All Features | #Drug Features | #Cancer features | | #Interactions | | #Nodes | | Performance ($R^2$) | | | |
|---|---|---|---|---|---|---|---|---|---|---|---|---|---|
| | | | | Residue | Others | DxM | Others | 1st | 2nd | NN | RF | SVM | Lasso |
| AG | 2,971 | 62 | 1 | 6 | 20 | 5 | 30 | 38 | 8 | 0.031 | 0.026 | 0.026 | **0.049** |
| Bone | 3,410 | 84 | 1 | 5 | 25 | 2 | 51 | 10 | 0 | 0.066 | 0.047 | 0.062 | **0.086** |
| Breast | 4,706 | 129 | 1 | 14 | 59 | 1 | 54 | 26 | 6 | 0.074 | 0.075 | 0.086 | **0.110** |
| CNS | 4,250 | 114 | 2 | 14 | 47 | 2 | 49 | 11 | 0 | 0.055 | 0.013 | 0.042 | **0.063** |
| Cervix | 1,044 | 37 | 1 | 9 | 15 | 1 | 11 | 7 | 0 | 0.103 | 0.132 | 0.144 | **0.165** |
| Endometrium | 1,073 | 33 | 1 | 3 | 25 | 0 | 4 | 11 | 4 | **0.118** | 0.045 | 0.049 | 0.064 |
| Haematopoietic | 4,204 | 119 | 2 | 14 | 48 | 12 | 43 | 11 | 0 | **0.138** | 0.098 | 0.093 | 0.127 |
| Kidney | 2,458 | 73 | 1 | 3 | 26 | 0 | 43 | 9 | 0 | 0.084 | 0.063 | 0.068 | **0.092** |
| Large intestine | 4,628 | 141 | 0 | 32 | 69 | 9 | 31 | 12 | 0 | **0.120** | 0.087 | 0.091 | 0.112 |
| Liver | 1,348 | 48 | 0 | 7 | 18 | 0 | 23 | 7 | 0 | 0.031 | 0.067 | 0.067 | **0.086** |
| Lung (NSCLC) | 9,205 | 207 | 0 | 31 | 79 | 12 | 85 | 15 | 0 | **0.114** | 0.086 | 0.085 | 0.109 |
| Lung (others) | 7,206 | 162 | 0 | 29 | 58 | 11 | 64 | 30 | 6 | **0.120** | 0.078 | 0.075 | 0.101 |
| Lymphoid | 13,302 | 291 | 2 | 41 | 99 | 17 | 132 | 18 | 0 | 0.078 | 0.091 | 0.086 | **0.108** |
| Oesophagus | 3,337 | 91 | 1 | 10 | 46 | 0 | 34 | 10 | 0 | 0.044 | 0.022 | 0.030 | **0.054** |
| Ovary | 3,502 | 113 | 1 | 16 | 65 | 2 | 29 | 11 | 0 | **0.107** | 0.058 | 0.063 | 0.088 |
| Pancreas | 2,421 | 84 | 1 | 11 | 24 | 0 | 48 | 10 | 0 | 0.057 | 0.077 | 0.072 | **0.095** |
| Pleura | 1,431 | 36 | 0 | 5 | 11 | 2 | 18 | 11 | 4 | 0.072 | 0.066 | 0.063 | **0.089** |
| Skin | 5,732 | 132 | 1 | 17 | 62 | 3 | 49 | 12 | 0 | 0.033 | 0.021 | 0.041 | **0.062** |
| Soft tissue | 1,938 | 63 | 1 | 6 | 34 | 2 | 20 | 8 | 0 | **0.106** | 0.069 | 0.073 | 0.103 |
| Stomach | 2,327 | 83 | 1 | 15 | 49 | 1 | 17 | 20 | 5 | 0.055 | 0.048 | 0.048 | **0.067** |
| Thyroid | 1,352 | 33 | 1 | 2 | 14 | 0 | 16 | 6 | 0 | 0.058 | 0.065 | 0.081 | **0.108** |
| UAT | 3,856 | 126 | 2 | 19 | 47 | 6 | 52 | 12 | 0 | 0.051 | 0.022 | 0.046 | **0.072** |
| Urinary tract | 1,454 | 68 | 2 | 5 | 28 | 1 | 32 | 9 | 0 | 0.051 | 0.039 | 0.052 | **0.072** |
| Overall | 87,155 | | | | | | | | | **0.125** | 0.103 | 0.086 | 0.116 |

The best performance for each cancer type is highlighted in bold. AG: autonomic ganglia; CNS: central nervous system; DxM: drug-mutation interaction term; NN: neural networks; NSCLC: non-small cell lung cancer; $R^2$: coefficient of determination; RF: random forests; SVM: support vector machine; UAT: upper aerodigestive tract; $\#IC_{50}$: the number of drug responses; #Nodes: the number of nodes in the first and second hidden layers of neural networks.

**S6 Table . Prediction performances of using random 10X feature selection.**

| Cancer type | #$IC_{50}$ | #All Features | #Drug Features | #Cancer features | | #Interactions | | #Nodes | | Performance ($R^2$) | | | |
|---|---|---|---|---|---|---|---|---|---|---|---|---|---|
| | | | | Residue | Others | DxM | Others | 1st | 2nd | NN | RF | SVM | Lasso |
| AG | 2,971 | 620 | 7 | 71 | 220 | 49 | 273 | 38 | 8 | 0.052 | 0.005 | 0.040 | **0.075** |
| Bone | 3,410 | 840 | 11 | 63 | 255 | 31 | 480 | 10 | 0 | **0.135** | 0.017 | 0.087 | 0.126 |
| Breast | 4,706 | 1,290 | 14 | 183 | 533 | 45 | 515 | 26 | 6 | **0.496** | 0.046 | 0.147 | 0.198 |
| CNS | 4,250 | 1,140 | 15 | 172 | 434 | 22 | 497 | 11 | 0 | **0.551** | 0.040 | 0.135 | 0.170 |
| Cervix | 1,044 | 370 | 13 | 62 | 174 | 7 | 114 | 7 | 0 | **0.449** | 0.070 | 0.163 | 0.251 |
| Endometrium | 1,073 | 330 | 4 | 62 | 198 | 9 | 57 | 11 | 4 | **0.148** | 0.046 | 0.056 | 0.093 |
| Haematopoietic | 4,204 | 1,190 | 15 | 144 | 462 | 121 | 448 | 11 | 0 | **0.421** | 0.027 | 0.122 | 0.180 |
| Kidney | 2,458 | 730 | 15 | 68 | 265 | 0 | 382 | 9 | 0 | **0.483** | 0.045 | 0.180 | 0.221 |
| Large intestine | 4,628 | 1,410 | 7 | 348 | 672 | 87 | 296 | 12 | 0 | **0.289** | 0.045 | 0.100 | 0.174 |
| Liver | 1,348 | 480 | 17 | 67 | 174 | 19 | 203 | 7 | 0 | **0.566** | 0.041 | 0.090 | 0.148 |
| Lung (NSCLC) | 9,205 | 2,070 | 7 | 309 | 708 | 87 | 959 | 15 | 0 | 0.131 | 0.056 | 0.092 | **0.141** |
| Lung (others) | 7,206 | 1,620 | 10 | 215 | 546 | 68 | 781 | 30 | 6 | **0.389** | 0.059 | 0.113 | 0.164 |
| Lymphoid | 13,302 | 2,910 | 15 | 461 | 917 | 105 | 1,412 | 18 | 0 | **0.430** | 0.042 | 0.143 | 0.180 |
| Oesophagus | 3,337 | 910 | 9 | 115 | 426 | 20 | 340 | 10 | 0 | **0.361** | 0.022 | 0.063 | 0.096 |
| Ovary | 3,502 | 1,130 | 12 | 159 | 492 | 31 | 436 | 11 | 0 | **0.322** | 0.029 | 0.089 | 0.127 |
| Pancreas | 2,421 | 840 | 19 | 108 | 307 | 8 | 398 | 10 | 0 | **0.442** | 0.022 | 0.146 | 0.190 |
| Pleura | 1,431 | 360 | 8 | 46 | 132 | 16 | 158 | 11 | 4 | **0.440** | 0.044 | 0.117 | 0.158 |
| Skin | 5,732 | 1,320 | 15 | 174 | 555 | 49 | 527 | 12 | 0 | **0.388** | 0.006 | 0.082 | 0.126 |
| Soft tissue | 1,938 | 630 | 10 | 79 | 292 | 33 | 216 | 8 | 0 | **0.408** | 0.072 | 0.128 | 0.189 |
| Stomach | 2,327 | 830 | 15 | 141 | 388 | 32 | 254 | 20 | 5 | **0.212** | 0.003 | 0.073 | 0.124 |
| Thyroid | 1,352 | 330 | 15 | 29 | 130 | 8 | 148 | 6 | 0 | **0.403** | 0.088 | 0.199 | 0.255 |
| UAT | 3,856 | 1,260 | 22 | 161 | 445 | 54 | 578 | 12 | 0 | **0.329** | 0.006 | 0.117 | 0.164 |
| Urinary tract | 1,454 | 680 | 23 | 71 | 304 | 15 | 267 | 9 | 0 | **0.707** | 0.090 | 0.121 | 0.197 |
| Overall | 87,155 | | | | | | | | | **0.378** | -0.021 | 0.130 | 0.157 |

The best performance for each cancer type is highlighted in bold. AG: autonomic ganglia; CNS: central nervous system; DxM: drug-mutation interaction term; NN: neural networks; NSCLC: non-small cell lung cancer; $R^2$: coefficient of determination; RF: random forests; SVM: support vector machine; UAT: upper aerodigestive tract; #$IC_{50}$: the number of drug responses; #Nodes: the number of nodes in the first and second hidden layers of neural networks.

**S7 Table . Pathway enrichment analysis.**

| PANTHER pathway | Reference list | Observed | Expected | Fold enrichment | P-value | FDR |
|---|---|---|---|---|---|---|
| Angiogenesis | 173 | 6 | 0.38 | 15.83 | 2.46E-06 | 2.02E-04 |
| Ras Pathway | 74 | 4 | 0.16 | 24.67 | 2.53E-05 | 8.31E-04 |
| Inflammation mediated by chemokine and cytokine signaling pathway | 260 | 6 | 0.57 | 10.53 | 2.36E-05 | 9.69E-04 |
| PDGF signaling pathway | 148 | 5 | 0.32 | 15.42 | 2.05E-05 | 1.12E-03 |
| Wnt signaling pathway | 312 | 5 | 0.68 | 7.31 | 6.21E-04 | 1.70E-02 |
| JAK/STAT signaling pathway | 17 | 2 | 0.04 | 53.7 | 7.81E-04 | 1.83E-02 |
| Cytoskeletal regulation by Rho GTPase | 87 | 3 | 0.19 | 15.74 | 1.01E-03 | 2.06E-02 |
| Axon guidance mediated by Slit/Robo | 26 | 2 | 0.06 | 35.11 | 1.70E-03 | 3.11E-02 |
| Interferon-gamma signaling pathway | 29 | 2 | 0.06 | 31.48 | 2.09E-03 | 3.42E-02 |
| Apoptosis signaling pathway | 118 | 3 | 0.26 | 11.6 | 2.35E-03 | 3.51E-02 |
| EGF receptor signaling pathway | 134 | 3 | 0.29 | 10.22 | 3.34E-03 | 4.57E-02 |

**S8 Table . Drug-mutation relationships and their impact on $IC_{50}$ in NSCLC cells.**

| Interaction term | $IC_{50}$ impact | $\lvert IC_{50}$ impact$\rvert$ |
|---|---|---|
| PKA_102_CSV_X_Fingerprint_714 | -1.8652 | 1.8652 |
| PKA_260_HYD_X_Fingerprint_819 | -1.5855 | 1.5855 |
| PKA_247_HYD_X_Fingerprint_685 | 1.0754 | 1.0754 |
| PKA_200_HYD_X_Fingerprint_673 | 0.7291 | 0.7291 |
| PKA_197_B62_X_Fingerprint_576 | 0.5091 | 0.5091 |
| **PKA_187_CHA_X_Fingerprint_791** | **-0.4563** | **0.4563** |
| PKA_112_POL_X_Fingerprint_659 | -0.4440 | 0.4440 |
| PKA_244_ENG_X_Fingerprint_576 | 0.3811 | 0.3811 |
| PKA_73_ENG_X_Fingerprint_611 | 0.3802 | 0.3802 |
| PKA_73_POL_X_Fingerprint_611 | 0.3772 | 0.3772 |
| PKA_187_POL_X_Fingerprint_791 | 0.3621 | 0.3621 |
| PKA_226_HYD_X_Fingerprint_576 | 0.3613 | 0.3613 |
| PKA_293_X_Fingerprint_611 | 0.2765 | 0.2765 |
| PKA_187_B62_X_Fingerprint_826 | 0.2702 | 0.2702 |
| PKA_293_X_Fingerprint_647 | -0.2575 | 0.2575 |
| PKA_229_EXP_X_Fingerprint_576 | 0.2542 | 0.2542 |
| PKA_197_EXP_X_Fingerprint_576 | 0.2540 | 0.2540 |
| PKA_142_X_Fingerprint_611 | -0.2385 | 0.2385 |
| PKA_229_HYD_X_Fingerprint_576 | 0.2296 | 0.2296 |
| PKA_270_POL_X_Fingerprint_576 | 0.2009 | 0.2009 |
| PKA_270_HYD_X_Fingerprint_611 | 0.1979 | 0.1979 |
| PKA_260_POL_X_Fingerprint_819 | 0.1119 | 0.1119 |
| PKA_283_POL_X_Fingerprint_647 | -0.1099 | 0.1099 |
| PKA_280_ENG_X_Fingerprint_646 | 0.1027 | 0.1027 |
| PKA_226_X_Fingerprint_644 | 0.0963 | 0.0963 |
| PKA_293_EXP_X_Fingerprint_363 | -0.0886 | 0.0886 |
| PKA_73_ENG_X_Fingerprint_644 | 0.0852 | 0.0852 |
| PKA_216_ASA_X_Fingerprint_646 | -0.0618 | 0.0618 |
| PKA_73_EXP_X_Fingerprint_702 | -0.0443 | 0.0443 |
| PKA_102_VOL_X_Fingerprint_714 | -0.0433 | 0.0433 |
| PKA_283_POL_X_Fingerprint_644 | -0.0403 | 0.0403 |
| PKA_160_HYD_X_Fingerprint_696 | -0.0346 | 0.0346 |
| PKA_175_ENG_X_Fingerprint_685 | 0.0342 | 0.0342 |
| PKA_270_EXP_X_Fingerprint_611 | 0.0276 | 0.0276 |
| PKA_252_ASA_X_Fingerprint_646 | -0.0227 | 0.0227 |
| PKA_283_ASA_X_Fingerprint_576 | 0.0202 | 0.0202 |
| PKA_187_ASA_X_Fingerprint_791 | -0.0192 | 0.0192 |
| PKA_283_ASA_X_Fingerprint_647 | 0.0119 | 0.0119 |
| PKA_197_VOL_X_Fingerprint_702 | 0.0118 | 0.0118 |
| PKA_197_ASA_X_Fingerprint_798 | 0.0097 | 0.0097 |
| **PKA_187_VOL_X_Fingerprint_826** | **-0.0090** | **0.0090** |
| PKA_123_VOL_X_Fingerprint_363 | -0.0079 | 0.0079 |
| PKA_77_ASA_X_Fingerprint_714 | -0.0055 | 0.0055 |
| PKA_73_CHA_X_Fingerprint_714 | -0.0027 | 0.0027 |
| PKA_283_VOL_X_Fingerprint_673 | -0.0024 | 0.0024 |
| PKA_283_ASA_X_Fingerprint_644 | -0.0023 | 0.0023 |
| PKA_270_VOL_X_Fingerprint_673 | -0.0004 | 0.0004 |

The features illustrated in Fig 4 are highlighted in bold.

**S9 Table . Cancer cell line features.**

| Feature level | Feature | Nomenclature | Value |
|---|---|---|---|
| Residue | PKA position | PKA_[POSITION] | $x_i = \sum_{k=1}^{K} M_{ki}\omega, \omega = \{1, CSV_{ki}, EXP_k\}$ |
| | Mutant type | PKA_[POSITION]_[MT] | $x_{im} = \sum_{k=1}^{K} M_{kim}\omega, \omega = \{1, CSV_{ki}, EXP_k\}$ |
| | Charge | PKA_[POSITION]_CHA | $x_i = \sum_{k=1}^{K} C_{ki}$ |
| | Polarity | PKA_[POSITION]_POL | $x_i = \sum_{k=1}^{K} P_{ki}$ |
| | Hydrophobicity | PKA_[POSITION]_HYD | $x_i = \sum_{k=1}^{K} H_{ki}$ |
| | Accessible surface area | PKA_[POSITION]_ASA | $x_i = \sum_{k=1}^{K} A_{ki}$ |
| | Side-chain volume | PKA_[POSITION]_VOL | $x_i = \sum_{k=1}^{K} V_{ki}$ |
| | Energy per residue | PKA_[POSITION]_ENG | $x_i = \sum_{k=1}^{K} E_{ki}$ |
| | Substitution score | PKA_[POSITION]_B62 | $x_i = \sum_{k=1}^{K} S_{ki}$ |
| Motif | Sequence motif | MOT_2D_[NAME] | $x_t = \sum_{k=1}^{K}\sum_{n=1}^{N_k} M_{kn}L_t(k,n)\omega, \omega = \{1, CSV_{kn}, EXP_k\}$ |
| | Structural motif | MOT_3D_[NAME] | $x_T = \sum_{k=1}^{K}\sum_{n=1}^{N_k} M_{kn}L_T(k,n)\omega, \omega = \{1, CSV_{kn}, EXP_k\}$ |
| Domain | Subdomain | SDOM_[NAME] | $x_d = \sum_{k=1}^{K}\sum_{n=1}^{N_k} M_{kn}L_d(k,n)\omega, \omega = \{1, CSV_{kn}, EXP_k\}$ |
| | Functional domain | DOM_[NAME] | $x_D = \sum_{k=1}^{K}\sum_{n=1}^{N_k} M_{kn}L_D(k,n)\omega, \omega = \{1, CSV_{kn}, EXP_k\}$ |
| Gene | Mutation | MUT_[GENE] | $x_k = M_k\omega = \sum_{n=1}^{N_k} M_{kn}\omega, \omega = \{1, CSV_{kn}, EXP_k\}$ |
| | Expression | EXP_[GENE] | $x_k = EXP_k$, from GDSC |
| | Copy number variation | CNV_[GENE] | $x_k = CNV_k = \{gain, neutral, loss\}$, from COSMIC |
| Family | Family | SFAM_[NAME] | $x_f = \sum_{k=1}^{K} M_k F_f(k)\omega = \sum_{k=1}^{K}\sum_{n=1}^{N_k} M_{kn}F_f(k)\omega, \omega = \{1, CSV_{kn}, EXP_k\}$ |
| | Group | FAM_[NAME] | $x_g = \sum_{k=1}^{K} M_k G_g(k)\omega = \sum_{k=1}^{K}\sum_{n=1}^{N_k} M_{kn}G_g(k)\omega, \omega = \{1, CSV_{kn}, EXP_k\}$ |
| Pathway | Reaction | REC_[REACTOME_ID] | $x_r = \sum_{k=1}^{K} M_k R_r(k)\omega = \sum_{k=1}^{K}\sum_{n=1}^{N_k} M_{kn}R_r(k)\omega, \omega = \{1, CSV_{kn}, EXP_k\}$ |
| | Pathway | PWY_[REACTOME_ID] | $x_w = \sum_{k=1}^{K} M_k W_w(k)\omega = \sum_{k=1}^{K}\sum_{n=1}^{N_k} M_{kn}W_w(k)\omega, \omega = \{1, CSV_{kn}, EXP_k\}$ |
| | Biological process | GO_[GO_ID] | $x_b = \sum_{k=1}^{K} M_k B_b(k)\omega = \sum_{k=1}^{K}\sum_{n=1}^{N_k} M_{kn}B_b(k)\omega, \omega = \{1, CSV_{kn}, EXP_k\}$ |
| Sample | Primary site | CLS_Primary_site | From COSMIC |
| | Site subtype 1 | CLS_Site_subtype_1 | |
| | Site subtype 2 | CLS_Site_subtype_2 | |
| | Site subtype 3 | CLS_Site_subtype_3 | |
| | Primary histology | CLS_Primary_histology | |
| | Histology subtype 1 | CLS_Histology_subtype_1 | |
| | Histology subtype 2 | CLS_Histology_subtype_2 | |
| | Histology subtype 3 | CLS_Histology_subtype_3 | |
| | Microsatellite instability | CLS_msi | |
| | Average ploidy | CLS_average_ploidy | |
| | Tumour source | CLS_tumour_source | |
| | Age | CLS_age | |
| | Gender | CLS_gender | |
| | NCI code | CLS_NCI_code | |

$M_{ki}$: if the residue of protein kinase $k$ aligned to PKA position $i$ is mutated (1) or not (0); $CSV_{ki}$: the conservation score of the residue of protein kinase $k$ aligned to PKA position $i$; $EXP_k$: the gene expression level of protein kinase $k$; $M_{kim}$: if the residue of protein kinase $k$ aligned to PKA position $i$ is mutated to the amino acid type $m$ (1) or not (0); $C_{ki}$, $P_{ki}$, $H_{ki}$, $A_{ki}$, $V_{ki}$, or $E_{ki}$: respectively mean the charge, polarity, hydrophobicity, accessible surface area, side-chain volume, or energy differences caused by the mutated residue of protein kinase $k$ aligned to PKA position $i$; $S_{ki}$: the BLOSUM62 substitution score of the mutated residue of protein kinase $k$ aligned to PKA position $i$; $N_k$: the length of protein kinase $k$ sequence; $M_{kn}$: if the $n$th residue of protein kinase $k$ is mutated (1) or not (0); $L_t(k,n)$, $L_T(k,n)$, $L_d(k,n)$, or $L_D(k,n)$: respectively mean if the $n$th residue of protein kinase $k$ is located in sequence motif $t$, structural motif $T$, subdomain $d$, or functional domain $D$ (1) or not (0); $CSV_{kn}$: the conservation score of the $n$th residue of protein kinase $k$; $CNV_k$: the copy number variation status of protein kinase $k$; $F_f(k)$ or $G_g(k)$: respectively mean if protein kinase $k$ belongs to family $f$ or group $p$ (1) or not (0); $R_r(k)$, $W_w(k)$, or $B_b(k)$: respectively mean if protein kinase $k$ is implicated in reaction $r$, pathway $w$, or biological process $b$ (1) or not (0); NCI code: National Cancer Institute (NCI) Thesaurus code.
